# Guiding eQTL mapping and genomic prediction of gene expression in three pig breeds with tissue-specific epigenetic annotations from early development

**DOI:** 10.1101/2025.03.05.641618

**Authors:** Fanny Mollandin, Hervé Acloque, Maria Ballester, Marco Bink, Mario Calus, Daniel Crespo-Piazuelo, Pascal Croiseau, Sarah Djebali, Sylvain Foissac, Hélène Gilbert, Elisabetta Giuffra, Cervin Guyomar, Ole Madsen, Marie-José Mercat, Bruno da Costa Perez, Jani de Vos, Andrea Rau

## Abstract

Gene expression is a dynamic phenotype influenced by tissue-specific regulatory mechanisms, which can modulate expression directly or indirectly through cis or trans factors. Identifying genetic variants in these regulatory regions can improve both expression quantitative trait locus (eQTL) mapping and gene expression prediction. Whole genome sequences offer the possibility for enhanced eQTL mapping accuracy, but detecting causal variants remains challenging. Here, we evaluate the potential added-value of integrating tissue-specific epigenetic annotations, such as chromatin accessibility and methylation status, into within-breed genomic predictions of expression for three pig breeds. Functional annotations from early developmental stages improved eQTL mapping interpretability as shown by the enrichment of trait-relevant QTLs. However, despite the use of functional annotations, predictions across breeds remain challenging due to differences in genetic architectures. Our work contributes to the understanding of gene expression regulation in livestock and highlights the value of functional annotations, despite continued challenges for predictions across breeds.

**Highlights:**

- Matched whole-genome sequences and expressions reveal transcriptional regulatory variants.
- Early epigenetic marks can guide transcriptome prediction and eQTL mapping.
- Annotation-guided models can enhance the interpretability of eQTL mapping.
- Porting predictions across pig breeds remains challenging even with annotations.
- A case study linked *IGF2* liver eQTLs with annotations and cholesterol QTLs.

## 1. Introduction

Gene expression represents a heritable intermediate phenotype that can be expected to be more closely tied to the genome than conventional phenotypes (Stranger et al. 2007). Expression is notably impacted by tissue-specific regulatory mechanisms, which may have properties of inhibition, modulation, or promotion; examples include promoters, enhancers or repressors, which are located in non-coding or more rarely coding genomic regions (Ong and Corces 2011). Regulatory sequences may modulate gene expression either directly (*cis* factors, generally in proximity to the gene on the same chromosome), or indirectly by acting as *trans* factors, located more distantly (Wittkopp, Haerum, and Clark 2004). These regulatory sequences can be affected by mutations, including for instance single nucleotide polymorphisms (SNPs), in turn impacting gene expression. Identifying and capitalizing on these genetic variants holds promise for improved expression quantitative trait locus (eQTL) mapping and prediction of gene expression based on genomic sequence (Agarwal and Shendure 2020; Bessière et al. 2018). Large-effect variants in regulatory sequences are typically identified using expression genome-wide association studies (eGWAS), while more subtle effects remain difficult to detect, for example due to epistatic effects and low allele frequency of causal variants, leading to low linkage disequilibrium with variants available on commercial SNP-chips and thus insufficient power.

Although the use of WGS offers the possibility to discover new variants, its increasingly high dimension reduces statistical power for variant mapping and genomic prediction. A potential strategy to better exploit WGS data may be the prioritization of certain regions of the genome through the use of functional annotations. In particular, integrating information on regulatory mechanisms in predictions of gene expression from WGS data could lead to prediction gains and shed insight on the underlying regulatory processes (Avsec et al. 2021), thus providing a better understanding of the biological processes driving complex traits and consolidating future genome annotations. Many types of models have been proposed for simultaneous QTL mapping and genomic prediction of complex traits, including BayesR (Erbe et al. 2012), which has been shown to be well-suited for traits with a small number of moderate to strong QTLs (Mollandin, Rau, and Croiseau 2021; Moser et al. 2015). A natural extension of this approach is the BayesRC model (MacLeod et al. 2016), which proposes the use of functional annotations to partition SNPs into categories that are independently modeled with the BayesR mixture model.

One limitation of BayesRC is the need for disjoint annotation categories; however, complex, large-scale functional annotations now cover multiple tissues, temporalities, and types of assays (Clark et al. 2020), generating substantial overlaps across annotation categories. The recently proposed BayesRCπ model implemented in the BayesRCO software (Mollandin et al. 2022) overcomes this limitation by disambiguating among multiple annotations for SNPs by preferentially assigning them to the most representative annotation, thus making it a promising approach to account for complex, overlapping functional annotations. Current efforts to incorporate functional annotations into genomic prediction models have not generally led to significant gains in prediction accuracy for livestock production traits, although improvements in QTL mapping precision have been observed (Abdollahi-Arpanahi, Morota, and Peñagaricano 2017; Xiang et al. 2021). A practical framework for the most appropriate use of prior annotation information has yet to be identified, and the most relevant sets of annotations must typically be identified on a case-by-case basis.

Gene expression represents a dynamic phenotype that has been shown to be highly tissue-specific. Epigenetic marks, such as chromatin accessibility (Klemm, Shipony, and Greenleaf 2019) and methylation (Corbett et al. 2022), are particularly relevant for gene expression given the strong impact of epigenetic mechanisms on gene transcription. In early developmental stages, epigenetics determines cell fate which ultimately affects future organ function (Gerrard et al. 2020). This suggests the potential importance of using early developmental and tissue-specific epigenetic functional annotations to prioritize regions in tissue-specific eQTL analysis. Contrary to human and mouse, to date relatively few studies have identified regulatory elements and analyzed their phenotypic impact in livestock species. Recent efforts, such as the Functional Annotation of ANimal Genomes (FAANG) and Farm Animal Genotype-Tissue Expression (FarmGTEx) consortia, have sought to fill this gap by providing insight into the regulatory mechanisms of gene expression in multiple tissues and several livestock species, notably through rich publicly available catalogs of tissue-specific functional annotations in pigs and cattle (Clark et al. 2020; Liu et al. 2022; Teng et al. 2024). In this context, the GENE-SWitCH project (Acloque et al. 2022) generated extensive functional genomic annotations in a variety of different tissues during early developmental stages for both pig and chicken (https://data.faang.org/projects/GENE-SWitCH).

Among livestock species, pigs hold particular interest as both an important source of meat for humans and as a highly relevant human biomedical model due to their similarity in anatomical structure, physiology, and immunology (Lunney et al. 2021). Over the last century, pig breeding programs and the introduction of crosses with specialized dam and sire lines enabled rapid genetic improvements and the development of multiple breeds and more diverse breeding goals reflecting societal needs (Merks 2000; Neeteson-van Nieuwenhoven, Knap, and Avendaño 2013). For example, the Duroc (DU), Landrace (LD), and Large White (LW) commercial breeds differ considerably in muscle growth and structure (Lee et al. 2012; Tang et al. 2020). Previous eQTL studies in pigs have typically focused on breed-specific analyses (Ballester et al. 2017; Maroilley et al. 2017). A recent study instead sought to identify genetic polymorphisms associated with gene expression variability in the duodenum, liver, and muscle shared across the DU, LD and LW breeds, yielding a set of nearly 14 million significant *cis* and *trans* expression-associated regulatory variants within and across tissues (Crespo-Piazuelo et al. 2023).

In this work, we capitalize on the multi-breed and multi-tissue pig eQTL data from (Crespo-Piazuelo et al. 2023) to evaluate the potential added-value of tissue-specific epigenetic annotations for within-breed eQTL mapping and genomic prediction (subsequently referred to as annotation-guided eQTL mapping and annotation-guided prediction, respectively) of liver and muscle gene expression using base pair-resolution genotypes in three highly distinct commercial pig breeds: DU, LD and LW. In particular, we make use of tissue-specific annotations that were generated by the GENE-SWitCH project to characterize chromatin accessibility and methylation status at three different early developmental stages, and we focus on the expression of a target subset of genes of interest. Our study further evaluates the potential for these epigenetic annotations to prioritize shared regulatory mechanisms across breeds.

## 2. Results

### 2.1 Exploring the pertinence of tissue-specific epigenetic annotations for an intermediate molecular phenotype

Our objective was to leverage epigenetic annotations obtained at several early developmental stages, as well as predicted variant effects, to prioritize putative regulatory variants in genomic prediction models of tissue-specific gene expression in three commercial pig breeds (Figure 1). Transcriptome data in liver and muscle and genotype data were collected for a total of *n*=100 animals for each breed (DU, LD, LW). Rather than performing a transcriptome-wide analysis, we focused our attention on liver and muscle expression for a targeted subset of 10 genes distributed on 8 chromosomes (Supplementary Table 1). These included a set of 3 genes previously highlighted as being regulated by methylation: *IGF2* (Van Laere et al. 2003), *PRKAG1* (Kai et al. 2022) and *LEPR* (Hao, Cui, and Gu 2016). In addition, to explore the potential added-value of tissue-specific functional annotations for tissue-specific eQTL mapping and prediction, we focused on a subset of 7 genes (*DET1*, *NUDT22*, *SUPT3H*, *CELF2*, *R3HCC1*, *HUS1, SLA-7*) with *cis*-regulatory variants consistently found across multiple tissues (Crespo-Piazuelo et al. 2023). We hypothesize that leveraging tissue-specific annotations can reveal pertinent regulatory mechanisms for these genes beyond the previously identified shared signals across tissues.

**Figure 1.**
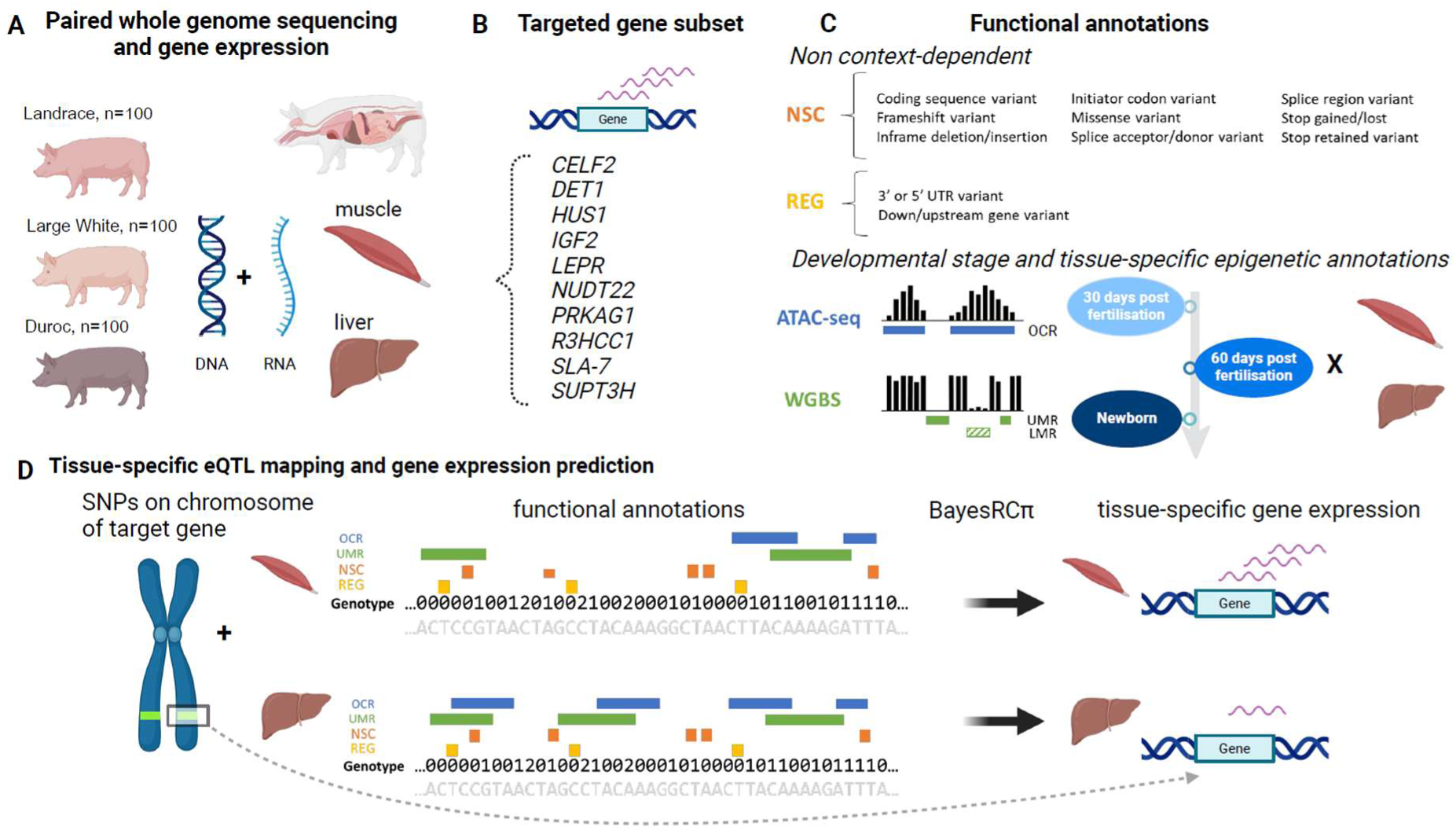
Schematic overview of the study. (A) Matched whole genome sequencing and liver and muscle gene expression data were available for *n*=100 pigs in each of three breeds (DU, LD, LW). (B) We focused on a targeted subset of 10 genes either regulated by methylation or with *cis*-regulatory variants found consistently across multiple tissues. (C) Functional annotations included both non-context dependent annotations and epigenetic annotations dependent on developmental stage and tissue. The former correspond to predicted variant effects (McLaren et al. 2016), i.e. those predicted to cause a nonsynonymous coding change (NSC) and those predicted to have potential regulatory roles (REG). The latter correspond to open chromatin regions (OCR) identified by assay for transposase-accessible chromatin using sequencing (ATAC-seq), as well as unmethylated regions (UMR) and lowly methylated regions (LMR) identified by whole genome bisulfite sequencing (WGBS). Epigenetic annotations were available for two tissues (liver, muscle) at three developmental stages: 30 days post-fertilisation (dpf), 70 dpf, and newborn. (D) Tissue-specific eQTL mapping and gene expression prediction were performed using BayesRCπ for each combination of {gene x tissue x breed} using variants on the respective chromosome of each gene and the full set of functional annotations. Created in BioRender. Mollandin, F. (2025) https://BioRender.com/l37n199

We used annotations generated post-mortem from independent Large White samples at early organogenesis (30 days post-fertilisation; dpf) and late organogenesis (70 dpf) as well as in newborn (NB) piglets (Acloque et al. 2022). We focused on liver-and muscle-specific epigenetic annotations generated at each developmental stage for two different functional genomic assays: methylation profiling by whole genome bisulfite sequencing (WGBS) and chromatin accessibility profiling by assay for transposase-accessible chromatin with high-throughput sequencing (ATAC-seq). For the former, genomic regions were categorized as being unmethylated (UMR), roughly corresponding to promoters, or lowly methylated (LMR), corresponding to putative enhancers (Stadler et al. 2011). For the latter, genomic regions were categorized as open chromatin regions (OCR), potentially representing enhancers, promoters, repressors or insulators (Roadmap Epigenomics Consortium et al. 2015). We additionally incorporated two broad tissue-agnostic annotations based on the Variant Effect Predictor (VEP) tool (McLaren et al. 2016): non-synonymous coding (NSC) variants and potential regulatory (REG) variants. Finally, tissue-specific epigenetic annotations (OCR, LMR, UMR) at each of the 3 developmental stages (30 dpf, 70 dpf, newborn) were concatenated with tissue-agnostic predicted variant effect categories (REG, NSC), and SNPs were assigned to one or more annotation category according to their genomic position; any non-annotated variants were assigned to a final “other” category.

For each of the 10 targeted genes, we then used as learning data the *n*=100 animals within each breed as well as the set of genetic variants located on the respective chromosome of the gene. In this way, our genomic predictions were constructed using an extensive definition of *cis*-regulatory variants. Genomic prediction models that were agnostic to annotations (BayesR) or that incorporated the aforementioned annotations as prior information (BayesRCπ) were fit to each learning dataset to perform eQTL mapping. We subsequently evaluated the portability of prediction models across breeds by assessing the prediction quality of the learning model on each of the two remaining breeds.

### 2.2 Considerable genomic and transcriptomic variability is observed among three commercial pig breeds

After removing genetic variants with a minor allele frequency (MAF) < 5% or with >10% missing genotype calls in the full set of *n*=300 animals, the original genomic dataset included 25,315,878 polymorphisms (Crespo-Piazuelo et al. 2023); for the set of 8 chromosomes corresponding to the 10 target genes, the number of SNPs varied from 542,356 (chr18) to 1,815,427 (chr1). Given our objective of evaluating the portability of genomic prediction models across breeds, we applied an additional filter to remove variants with a per-breed MAF < 5% in one or more of the breeds considered, which removed approximately a third of these variants (Supplementary Table 2); a total of 14,489,226 polymorphisms were retained across the 8 chromosomes, varying from 177,240 (chr18) to 629,069 (chr1) per chromosome, that were relatively uniformly distributed across each chromosome (Supplementary Figure 1). After filtering, we observed considerable genomic (Figure 2A) and, to a lesser extent, transcriptomic (Figure 2B) heterogeneity among the three pig breeds likely due to differences resulting from selection or drift. As expected, a strong separation of DU from the two other breeds is seen along the first principal component, particularly for the genomic and muscle transcriptome data, perhaps reflecting stronger selection constraints on muscle than liver. A secondary separation between LD and LW is present on the second principal component, with a weaker distinction observed in the liver transcriptome. Similar separation between the three pig breeds is observed for chromosome-specific genomic principal components analyses (Supplementary Figure 2).

**Figure 2.**
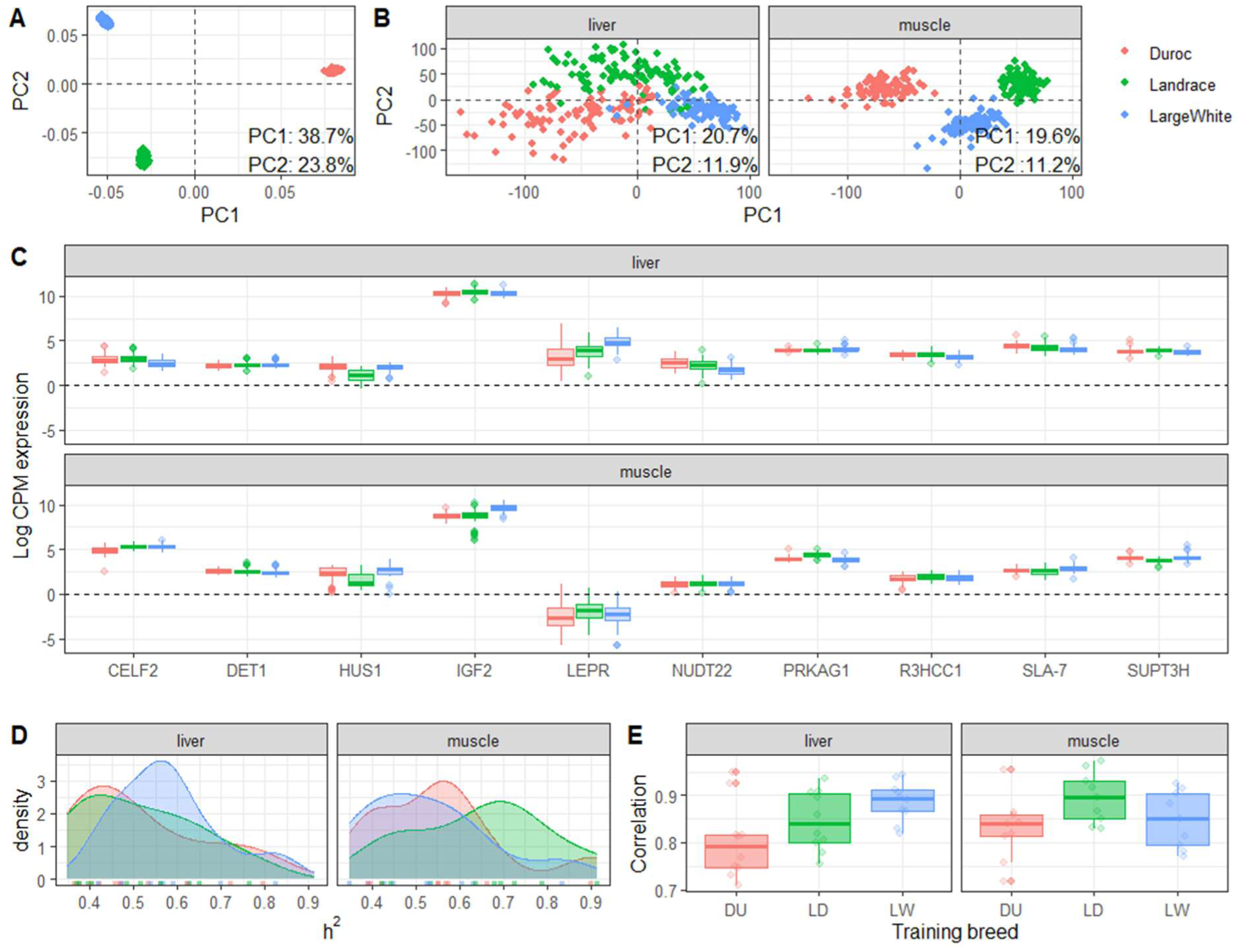
Genome-and transcriptome-wide heterogeneity among three pig breeds. (A) Principal components analysis (PCA) on genomic data from autosomal chromosomes. (B) PCA on transcriptomic data (sex-corrected log counts per million values; CPM) by tissue. Percentage variance explained for each of the first two components is indicated in text for each PCA. (C) Boxplots of transcriptomic data (sex-corrected log CPM values) for target genes by breed and tissue. (D) Distribution of estimated heritabilities of expression (across targeted genes and breeds) in liver and muscle, colored by the training breed. (E) Spearman correlations across targeted genes in learning data by breed and tissue. DU=Duroc (red); LD=Landrace (green); LW=Large White (blue).

As previously noted, our focus here is on the liver-and muscle-specific expression of 10 target genes (Figure 2C); note that as *LEPR* displays very low expression in the muscle, we consider only its expression in the liver in the following. Variability in the expression of these target genes is observed both across breeds within a given tissue (e.g., *LEPR* in liver) as well as between genes (e.g., generally strong expression for *IGF2*, generally weak expression for *NUDT22*). In the original study design, the DU and LD breeds included both males and females, whereas the LW breed had only males; to avoid sex-specific biases, log_2_ counts per million (CPM) expression values for the DU and LD breeds were corrected for an intercept (i.e., overall mean) and sex fixed effect, while those for the LW breed were corrected only for an intercept (Supplementary Figure 3).

Beyond the heterogeneity of the expression of these genes, we also observed considerable variability in the heritability of their expression, both between tissues as well as across breeds (Figure 2D). We further remark slightly higher expression heritabilities for LW in liver, and generally higher heritabilities in muscle for LD. These breed-specific differences are also reflected by disparities in the learning quality for each breed, as quantified by the correlation between observed and predicted expression values for each breed, with generally higher learning prediction accuracies for LD models trained on muscle compared to liver, and for LW models trained on liver compared to muscle (Figure 2E).

### 2.3 Annotations related to tissue-specific epigenetic marks during early development are biologically relevant for the analysis of complex traits

Annotations related to the predicted potential impact of variants as well as tissue-specific epigenetic marks such as chromatin accessibility and methylation are of particular interest for gene expression data, both with respect to prediction and for identifying potentially important regulatory regions. Overall, 16.9% and 14.1% of the filtered variants on the 8 chromosomes were assigned to one or more epigenetic or predicted variant effect categories in liver and muscle, respectively (Supplementary Figure 4A). Although most variants were assigned to a single annotation, a considerable number were assigned to multiple categories, with a maximum number of multi-annotations corresponding to 8 (Supplementary Figure 4B); generally similar patterns were observed between the two tissues. When considering each of the annotation categories separately (Supplementary Figure 4C, Supplementary Table 2), we remark that the two VEP categories are those with the sparsest (NSC) and densest (REG) densities. Tissue-and stage-specific epigenetic annotations were intermediate in terms of the number of annotated SNPs, with a slightly higher density in liver than muscle, particularly for early developmental stages (OCR at 30 dpf; LMR and UMR at 30 and 70 dpf, respectively). Annotation categories unsurprisingly tended to cluster together (Supplementary Figure 4D) first according to type of epigenetic mark (OCR, UMR, LMR) and predicted variant effect (NSC, REG), followed by tissue type (liver, muscle); we also generally remark a consistency between developmental stages and the annotation category clustering structure.

To evaluate the extent to which these annotations colocalized with previously identified variants of interest in GWAS of pig traits (Figure 3), we then tested each category for overrepresentation of curated QTLs for high-level trait categories (Meat and Carcass Traits, Health Traits, Exterior Traits, Production Traits and Reproduction Traits) identified in the PigQTLdb (Hu, Park, and Reecy 2021). The distribution of QTLs for each high-level trait category across chromosomes is shown in Supplementary Figure 5. Our results suggest that there is little overlap between LMR variants and previously identified QTLs in both liver and muscle; similarly, the variant effect prediction categories (REG, NSC) did not show strong enrichment. However, the remaining tissue-specific epigenetic categories were found to be highly enriched in PigQTLdb QTLs, supporting the biological relevance of these categories with respect to the genetic architecture of complex traits. A complementary analysis of the enrichment results of each high-level QTL trait category for annotations is shown in Supplementary Figure 6.

**Figure 3.**
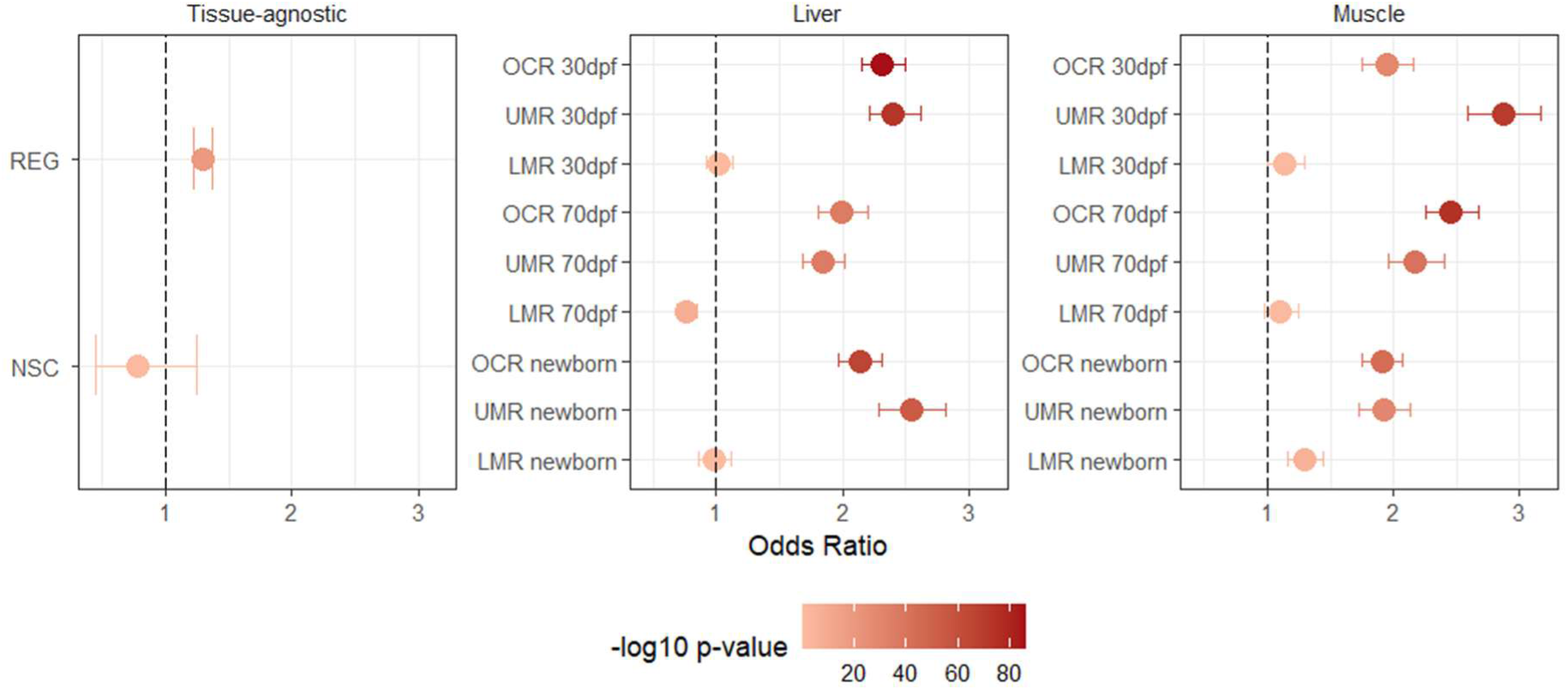
Annotations related to predicted variant effects and tissue-specific epigenetic marks during early development. Odds ratio and associated-log_10_ P-values from a Fisher’s exact test of the enrichment of QTLs from PigQTLdb broad trait categories within each grouping of annotation categories: tissue-agnostic VEP annotations (left), liver-specific (middle) and muscle specific (right) epigenetic annotations. OCR: open chromatin regions; UMR: unmethylated regions: LMR: lowly methylated regions; REG: predicted regulatory variant; NSC: predicted nonsynonymous coding variants; dpf: days post fertilization.

### 2.4 eQTL mapping is enriched in pig trait QTLs when guided by functional annotations in muscle and liver

We next sought to evaluate the impact of leveraging predicted variant effects and tissue-specific epigenetic annotations to prioritize variants in genomic prediction models fit for each breed independently. As previously noted, these annotations reflect partially redundant prior information (Supplementary Figure 4B), where SNPs may be simultaneously assigned to one or more categories. The annotation-agnostic BayesR genomic prediction model (Erbe et al. 2012), which is based here on a prior five-component mixture model (null, very small, small, medium or large SNP effects), was recently extended in the BayesRCπ model (Mollandin et al. 2022) by incorporating complex overlapping variant annotations through a prior distribution. For both BayesR and BayesRCπ, eQTL mapping can be performed by ranking SNPs according to their estimated posterior variance (see Material and Methods).

One measure of the biological pertinence of highly ranked SNPs is the enrichment of known QTLs among those with large estimated posterior variances (Supplementary Figure 7). When investigating the distribution of gene set enrichment analysis (GSEA) results for each PigQTLdb trait across the 10 target genes and 3 learning breeds for each tissue, we remark significantly stronger enrichment of known QTLs for the annotation-aware BayesRCπ model as compared to the annotation-agnostic BayesR model in both tissues (Figure 4A; Kolmogorov-Smirnov test, *P* = 2.5 x 10^-16^ and *P =* 2.3 x 10^-9^ for liver and muscle, respectively). When specifically considering the number of scenarios (10 target genes x 3 learning breeds x 5 PigQTLdb broad trait categories) where statistically significant enrichments were observed among highly ranked SNPs (Figure 4B), we remark that BayesRCπ generally leads to as many or more high-level trait categories with significant enrichments as compared to BayesR, with the exception of the Exterior traits in liver. In particular, in both tissues considerably more scenarios with significant QTL enrichments are observed for Reproduction and Meat and carcass traits when using annotations, and for Health traits in liver. Detailed results for each of the individual PigQTLdb traits with significant enrichment among estimated posterior variances are shown in Supplementary Tables 3 and 4 (BayesRCπ and BayesR, respectively).

**Figure 4.**
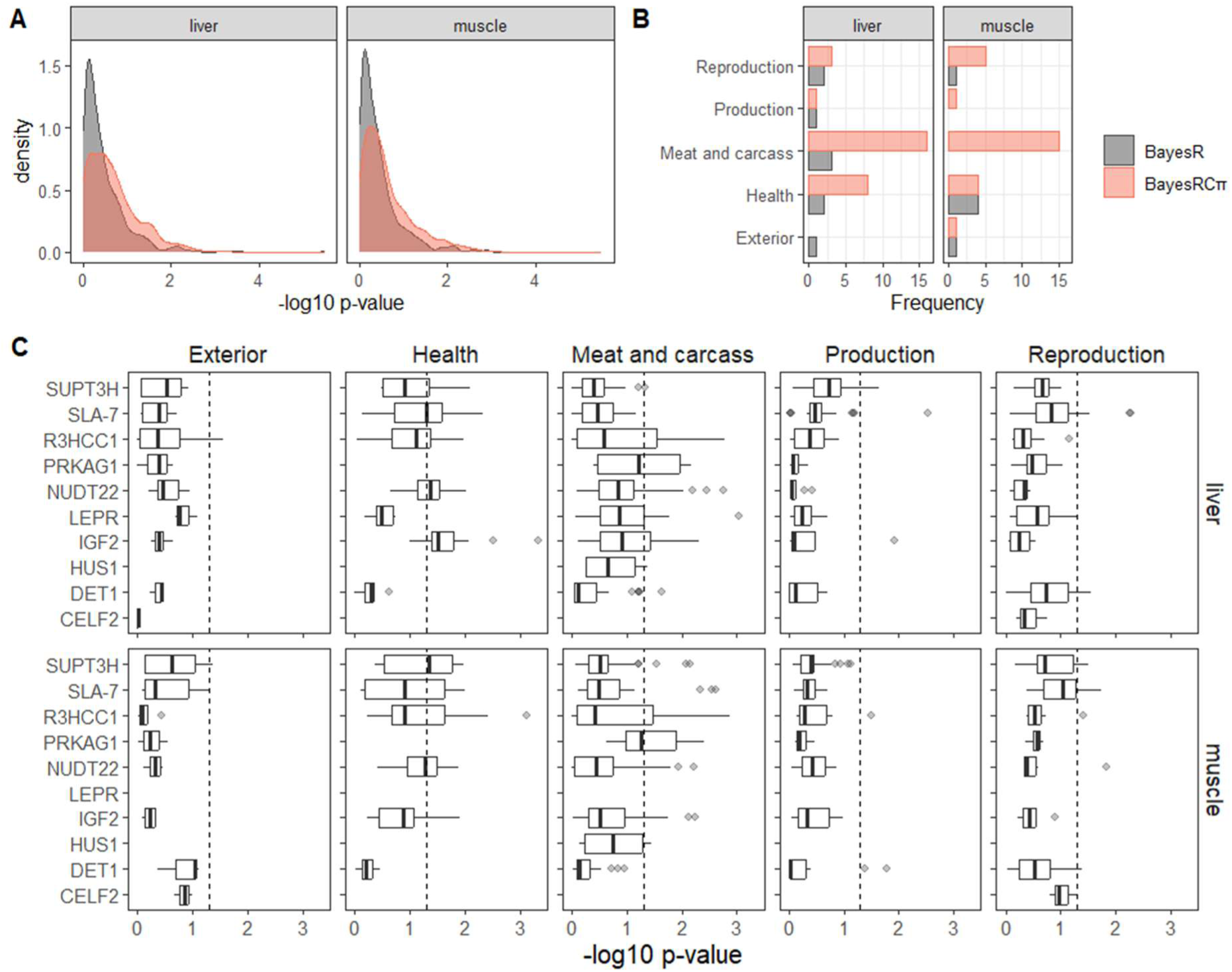
**Guiding eQTL mapping of gene expression in muscle and liver with functional annotations.**(A) Density plot of-log_10_ P-values for the enrichment of QTLs among estimated posterior variances for BayesR (grey) and BayesRCπ (orange) in each tissue across all scenarios (genes, learning breeds). (B) Number of scenarios (genes, learning breed) with significant enrichments (P-value < 0.05) in QTLs for different categories of pig traits based on estimated posterior variances for BayesR (grey) and BayesRCπ (orange). (C) Distribution (across learning breeds and individual traits) of-log_10_ P-values corresponding to the enrichment of QTLs for broad trait categories among BayesRCπ posterior variances for each gene within each tissue. In each panel, the dotted vertical line corresponds to a P-value threshold of 0.05 for reference.

We similarly investigated QTL enrichments among high-level trait categories in each tissue for each gene individually (Figure 4C). The distribution of enrichment P-values for BayesRCπ estimated posterior variances across scenarios (learning breeds x PigQTLdb trait within each high-level category) reveals an enrichment of Health QTLs among *IGF2* eQTLs in liver, as well as of Meat and carcass QTLs among *PRKAG1* eQTLs in both liver and muscle. Taken together, these results suggest that biologically relevant QTLs for a variety of trait categories are enriched in relevant tissues when using an annotation-guided model. We thus focus the remainder of this work on results from the BayesRCπ model.

### 2.5 Predicting expression across breeds is challenging, but not due to differences in pig trait QTL enrichments

Our results suggest that using predicted variant effects and tissue-specific epigenetic annotations leads to a more meaningful eQTL mapping with respect to known QTLs for a variety of trait categories (Figure 4). We next sought to evaluate whether these annotations were similarly useful for performing genomic predictions of gene expression across breeds. To assess the BayesRCπ prediction portability across breeds, we first learned model parameters on each breed individually and subsequently predicted gene expression values for the remaining two breeds (10 target genes x 3 learning breeds x 2 tissues, with the exception of *LEPR* with only liver-specific expression).

Globally, Spearman correlations between predicted and observed gene expression tended to be modest across genes for each learning-validation breed pair and quite variable across genes, with a maximum of 0.1561 on average (standard deviation = 0.228) when learning on LW and validating on LD in liver, and a minimum of 0.0173 (standard deviation = 0.240) on average when learning on LW and validating on DU in liver (Figure 5A). Given the strong genetic distinctness of DU compared to LD and LW (Figure 2A), it is somewhat surprising that these global trends do not align with the expectation of systematically worse performance for learning-validation breed pairs including DU.

**Figure 5.**
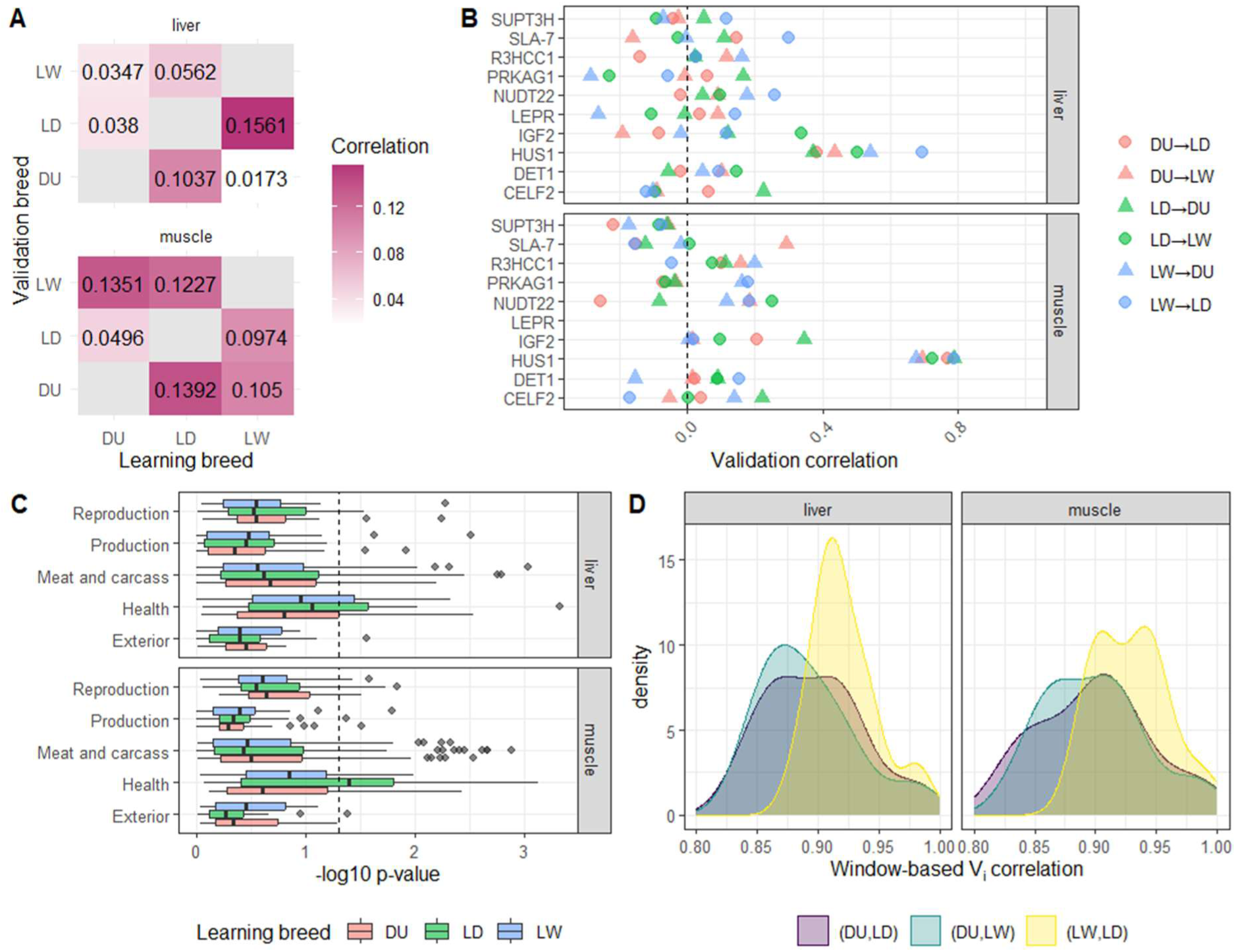
Predictions of expression in liver and muscle across breeds. (A) Heatmap of Spearman correlations from BayesRCπ for each combination of learning and validation breed within each tissue, averaged across genes. Grey boxes along the diagonal correspond to cases where the learning and validation data are the same. (B) Individual Spearman correlations by gene for each tissue and each combination of training and validation breed (indicated by different colors and shapes). The vertical dotted line indicates 0 as a reference point. (C) Boxplots (across genes and learning breeds) of-log_10_ P-values for the enrichment of broad trait QTLs per breed. DU=Duroc (red); LD=Landrace (green); LW=Large White (blue). (D) Distribution (across genes) of the correlations between window-based posterior variance estimates for each pair of learning breeds.

These global average trends do mask the considerable variability in prediction quality among genes for each tissue and among learning-validation breed pairs (Figure 5B). For example, *HUS1* has very strong predictive performance for across-breed predictions in both tissues (ranging from 0.373 to 0.791), in particular for muscle, regardless of the pair of breeds used for learning and validation. At the other extreme, *SUPT3H* in muscle has negative validation correlations (ranging from-0.219 to-0.058) for all combinations of learning and validation breeds. Genes with systematically negative correlations represent a particularly intriguing case. Focusing on top-ranked SNPs according to posterior variances (Supplementary Figure 8), greater genetic distinctness among breeds can be seen in a case with particularly weak prediction portability (e.g., *SUPT3H* in muscle, learning on DU) compared to one with high prediction portability (e.g., *HUS1* in muscle, learning on DU). Correlations between gene expression and the top PCs in these two cases are modest (Supplementary Table 5), but do show opposing trends for some PCs (e.g., a correlation of-0.215 and 0.165 for DU and LW, respectively, for PC1 of *SUPT3H*). Most genes are intermediate to these two cases, with no evident pattern revealing systematically advantageous or disadvantageous choices for learning and validation breed.

Expression predictions across breeds thus remain challenging despite the use of relevant annotations (Figure 3) as prior biological information, which is perhaps to be expected given the genetic and transcriptomic separation of these three breeds (Figure 2A and B) and the variability inherent in gene expression across breeds (Figure 2C and D). To investigate whether part of this difficulty could be explained by breed-specific PigQTLdb QTL enrichments among estimated posterior variances, we examined GSEA results across genes for each learning breed (Figure 5C). Although some variability can be observed across breeds, we do not observe any systematic breed-specific trends. Similarly, we sought to determine whether marked differences in inferred genetic architecture could play a role by investigating the distribution of correlations between sliding windows of cumulative estimated posterior variances for each pair of breeds (Figure 5D). Generally high correlations (> 0.90) were observed regardless of the pair of learning and validation breeds, although those including DU tended to be lower than that of LD and LW. Similar results were observed when estimating the correlation between individual rather than window-based estimated posterior variances (Supplementary Figure 9). Taken together, these results suggest that the difficulty of across breeds expression prediction is not linked to breed-specific annotation differences.

### 2.6 Annotation-driven eQTLs for IGF2 expression in liver colocalize with stage-specific epigenetic marks and cholesterol QTLs

Finally, we investigated in greater detail the role played by the predicted variant effects and tissue-specific epigenetic annotations in the eQTL mapping for a specific case study. Our previous results highlighted the strong enrichment of Health QTLs among BayesRCπ estimated posterior variances in liver for *IGF2*, which is located on chromosome 2 (Figure 4C). For *IGF2* liver expression, the Health trait with the most significant enrichment in both the DU and LD breeds (Supplementary Table 3) was LDL cholesterol (Chen et al. 2013); in the case of DU, a highly ranked SNP with a large posterior variance and previously identified as being associated with LDL cholesterol corresponded to chr2:50750725 (rs81359856; Figure 6A). The distribution of genotypes for this SNP in each breed (Figure 6B) unsurprisingly reveals an association with *IGF2* expression in DU, notably higher expression associated with the AA homozygote. A similar but less marked trend is seen for this SNP in the LD breed, whereas only a small frequency of AA homozygotes at this locus are observed in LW.

**Figure 6.**
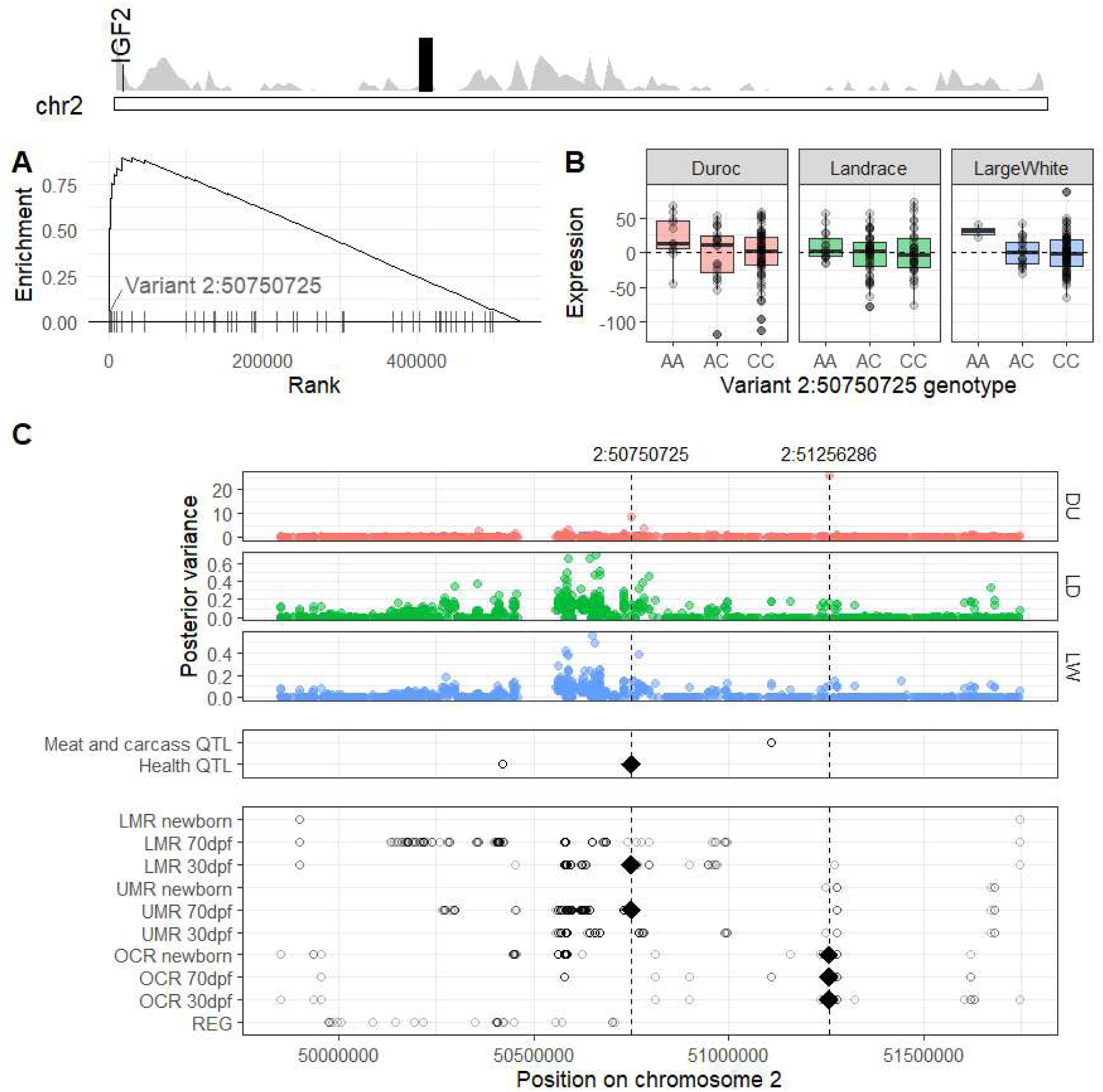
Annotation-guided eQTL mapping for *IGF2* expression in liver. (top) Position on chromosome 2 of *IGF2* and density of SNPs (grey) on chromosome 2, with the region visualized in panel C highlighted in black. (A) Enrichment plot for cholesterol QTLs from PigQTLdb based on posterior variances for liver *IGF2* expression in the DU breed. (B) Boxplots of top-ranked DU variant for cholesterol enrichment (chr2:50750725, panel A) in DU, LD and LW. (C) Neighborhood +/-1 Mb around variant chr2:50750725, a highly ranked variant for *IGF2* expression in liver. Estimated posterior variances of effects for each genetic variant in the window are shown in the middle panel for each learning breed (DU=Duroc: LD = Landrace; LW = Large White). The bottom panels indicate the position of QTL categories from PigQTLdb and of predicted variant effect annotations and liver-specific epigenetic annotations (circles). The position of variants chr2:50750725 and chr2:51256286 are indicated with a dotted line, and annotations located at these positions are highlighted with diamonds.

We next examined the colocalization of breed-specific eQTLs (based on estimated posterior variances) for *IGF2* liver expression with the predicted variant effect and tissue-specific annotations as well as the PigQTLdb high-level trait QTLs in a 1Mb window around this leading SNP (Figure 6C). Although some similarities can be seen in the genetic architectures identified for each breed from annotation-guided genomic prediction, breed-specific regions are indeed evident despite the use of a common set of annotations, in line with the results shown in Figure 5D. Similar conclusions can be drawn from chromosome-wide Manhattan plots of the window-based cumulative posterior variances for *IGF2* in both liver (Supplementary Figure 10) and muscle (Supplementary Figure 11), as well as in the 1Mb window directly surrounding *IGF2*, rather than around the leading SNP from the LDL cholesterol enrichment analysis (Supplementary Figure 12). By investigating the physical positions of each annotation as well as those of known PigQTLdb QTLs, we remark interesting overlaps between mapped eQTLs within each breed; for example, variant chr2:50750725 (rs81359856), which had a large posterior variance in DU, corresponding to a known Health QTL (Figure 6C, middle panel), is also annotated as being LMR at 30 dpf as well as UMR at 70 dpf (Figure 6C, bottom panel). As another example, the top ranked SNP for *IGF2* liver expression in DU in this window (chr2:51256286; rs341162083) is annotated as OCR at all three developmental stages (Figure 6C). Overall, the proportion of medium-to large-effect variants identified within each annotation category (Supplementary Figure 13) supports the biological relevance of the NSC predicted variant effect category as well as the accessible and unmethylated regions in early development (30 and 70 dpf), with early development LMRs and predicted regulatory variants showing weaker enrichment in such effects; similar results are observed across the three breeds.

## 3. Discussion

In this work, we used predicted variant effects and tissue-specific epigenetic marks from early developmental stages as informative prior annotations to guide genomic prediction models of liver and muscle gene expression in three commercial pig breeds displaying considerable genomic and, to a lesser extent, transcriptomic heterogeneity (Figure 2). Whole genome sequencing and RNA sequencing data respectively provided high-resolution genotypes and an intermediate molecular phenotype to construct predictive models for each breed. We focused on a target subset of 10 genes based on a previous eQTL analysis (Crespo-Piazuelo et al. 2023) and prior knowledge of the pertinence of epigenetic regulatory mechanisms. Although the preselection of a subset of genes represents a pertinent case study to evaluate the potential added value of functional annotations for genomic prediction and eQTL mapping, we note that it does limit a transcriptome-wide generalization of our results.

Tissue-specific epigenetic annotations during early development (early and late organogenesis, newborn) were shown to be enriched in previously identified QTLs of complex traits, thus supporting the pertinent biological signal therein (Figure 3). A model directly incorporating these annotations was shown to improve interpretability of eQTL mapping, as measured by the over-enrichment of QTLs of high-level trait categories among variants with large estimated posterior variances, in comparison to prediction based on genomic sequence alone (Figure 4). However, annotation-guided prediction models did not systematically lead to strong performance when performing predictions across breeds. Our results suggest that the challenge of porting predictions across breeds here is more likely to be linked to differences in genetic architectures rather than disparities in the pertinence of annotations (Figure 5). Similar challenges have been observed for the prediction of gene expression across human populations (Bhattacharya et al. 2022; Keys et al. 2020). However, we note that pig populations have much stronger average genetic relationships between individuals and increased linkage disequilibrium compared to human populations (Wray et al. 2019).

Despite similarities in highly prioritized genomic regions across breeds, differences in linkage disequilibrium, allele frequency, and fixed mutations due to intensive selection or drift complicate prediction across breeds even when incorporating relevant annotations to better target potential causal mutations. For example, around *IGF2*, a haplotype segregating at low frequency in the 1977 LW ancestral population increased to high frequency, with a particularly strong increase in a 1.5 Mb segment at the end of the region corresponding to the location of the *IGF2* gene, where the haplotype was almost fixed (Boitard et al. 2023). We provided a more in-depth look at the eQTL mapping results for *IGF2* liver expression, which were strongly enriched for health trait QTLs and showed interesting overlap with early development epigenetic marks (Figure 6). *IGF2* liver expression represents a particularly interesting case study due to its established link with hepatic lipid homeostasis (Kessler et al. 2016; Lopez et al. 2018), and its effect on growth and fatness traits (Tortereau et al. 2011).

Our annotation-guided model enabled the identification of eQTLs that may be good candidates for regulating *IGF2* expression in different breeds within the same genomic region. Importantly, the integration of functional annotations from several time points during early embryonic and foetal life prioritized both functionally and biologically relevant variants, in which we could identify enrichment of health-related traits, including LDL cholesterol. These results suggest the interest in using developmental-related functional annotations to identify genetic variants in putative regulatory regions that affect complex traits in adults, a central point that we sought to address within the European GENE-SWitCH project, which is part of the FAANG initiative (https://www.gene-switch.eu/).

There are several limitations of this work that are worth noting. First, variability in gene expression is both genetically driven as well as linked to complex tissue-specific regulatory mechanisms that respond to environmental stimuli, highlighting the challenge of predicting gene expression based on genotypes; compounding this difficulty are the modest learning sample sizes (*n=*100) available for each breed. While eQTL studies generally focus on a 1Mb window around genes, *cis--*eQTL effects only partially recapitulate gene expression heritability (Wang et al. 2024). Capturing distal *trans*-eQTL effects can improve our understanding of gene expression, although this comes with a heavy statistical testing burden (Võsa et al. 2021). To balance biologically reasonable assumptions with computational constraints, we used as explanatory variables in our annotation-guided models chromosome-wide variants. In future studies, it could be of interest to either widen or narrow the focus of our eQTL-mapping: working whole genome-wide may identify long-range regulatory mechanisms, while exploring cis-eQTLs alone may help increase detection power.

In addition, preprocessing steps for functional annotation data as well as the genomic and transcriptomic data are particularly important for annotation-guided genomic prediction and eQTL mapping. We have chosen here to ignore rare variants and reduce the number of SNPs in the WGS data by filtering on per-breed minor allele frequency. This filtering aims to achieve adequate detection power and focus on a common set of SNPs across breeds, but it is quite stringent given the modest sample sizes in each breed. Although this strategy enables cross-breed comparability, one major limitation is that it ignores potentially important breed-specific variants, e.g. those that have been fixed due to selection in a subset of breeds. In future work, it could be interesting to consider alternative filtering thresholds that better balance these aspects.

Epigenetic annotations were obtained from a different set of LW animals with a limited number of replicates, which leaves the possibility that they do not completely reflect potential regulatory mechanisms of the DU and LD breeds or that some regions have greater within-breed variability than that captured here. However, we did not observe any marked benefit in using LW epigenetic annotations for prioritizing biologically meaningful variants for LW eQTL mapping as compared to the two other breeds (Figure 5C). In addition, only annotations from early developmental stages (30 and 70 dpf, newborn) were used, while the tissue-specific expression data used as an intermediate molecular phenotype were collected from 6-month-old animals. Predictive models based on gene expression data sampled concomitantly to phenotypes were recently shown to improve phenotypic predictions (Perez et al. 2022); similarly, it might be reasonable to assume that the most meaningful annotations would correspond to those obtained at the stage closest to the sampling time of the expression data. However, a more complicated picture emerges from our results. For example, newborn UMRs have the highest proportion of medium-and large-effect variants for *IGF2* liver expression (Supplementary Figure 13), closely followed by chromatin accessibility during early and late organogenesis.

For future work, although extending this study to the full transcriptome would likely be computationally over-demanding, it would be interesting to conduct a comparative analogous study for a complementary set of genes. Alternative strategies for constructing prior categorization of variants according to epigenetic marks could also be considered. One plausible hypothesis would be to consider a category of SNPs in regions characterized by changes in methylation or chromatin accessibility between developmental stages, *i.e.* so-called regulatory switches, which may represent dynamic regulation mechanisms involved in gene expression. In addition, other annotations could prove to be useful, including for example breed-and tissue-specific chromatin conformation and histone marks or eQTL signals from PigGTeX (Teng et al. 2024); we leave such investigations to future work.

## 4. Conclusions

Genomic prediction models are widely used for the prediction of complex traits or disease risk, and they can similarly be used to perform eQTL mapping and prediction of gene expression data. We sought to construct predictive models of liver and muscle gene expression in pigs using base pair-resolution genotypes for three commercial breeds. We investigated the added-value of augmenting our predictive models of gene expression with prior annotations of polymorphisms based on predicted variant effects and tissue-specific epigenetic marks from early development. Our results suggested the utility of these functional annotations for eQTL mapping, though they did not systematically lead to strong predictive results across breeds. We postulate that this is due to differences in genetic architectures between breeds rather than disparities in the pertinence of annotations.

## 5. Methods

### 5.1 Study design

Samples from three different tissues (duodenum, liver, and muscle) were collected at slaughter from *n*=300 approximately 6-month-old pigs of three different breeds (Duroc, DU; Landrace, LD; Large White, LW), with *n*=100 animals per breed (Crespo-Piazuelo et al. 2023). The DU and LD breeds included data for both sexes (DU, 50 castrated males and 50 females; LD, 39 entire males and 61 females), while LW included only entire males. Genomic DNA was extracted from blood (DU, LD) or liver (LW), and RNA was extracted from duodenum, liver, and muscle. DNA and RNA were then paired-end (2 x 150bp) sequenced using the Illumina NovaSeq6000 platform. In the current study, we focus in particular on data arising from the liver and muscle.

### 5.2 Whole genome sequencing data

Full details about DNA extraction, sequencing, mapping, and variant calling are available in (Crespo-Piazuelo et al. 2023). Briefly, sequencing reads were mapped to the *Sscrofa11.1* reference genome assembly with BWA-MEM (Li 2012) and genetic variant calling was performed with GATK (McKenna et al. 2010). Further data preprocessing steps were carried out using PLINK (v1.07) (Chang et al. 2015) to separate variants by chromosome and breed, and to remove variants with any missing calls, a minor allele frequency (MAF) less than 5% in one or more breeds (*n*=100 animals per breed), or fixed heterozygosity in one or more breeds. To ensure compatibility, minor allele encoding within breeds was specified to be that of the full dataset including all three breeds (*n*=300).

### 5.3 Transcriptomics data

As above, full details about RNA extraction, sequencing, mapping, and quantification are available in (Crespo-Piazuelo et al. 2023). Briefly, RNA sequences were mapped to the *Sscrofa11.1* reference genome assembly with STAR (Dobin et al. 2013). Gene expression was quantified by counting reads aligning to genes using RSEM (Li and Dewey 2011) and normalized using the trimmed mean of M-values (TMM) approach (Robinson and Oshlack 2010), yielding log_2_ counts per million (CPM) values (Robinson, McCarthy, and Smyth 2010). Lowly expressed genes (CPM values < 10/minimum library size in millions) were filtered from further analyses. To avoid sex-specific biases, transcriptomic data for the DU and LD breeds were subsequently corrected by retaining residuals from a linear regression of sex and an intercept on log_2_ CPM values; as LW animals were all male, their log_2_ CPM values were corrected only for an intercept. Finally, all corrected log_2_ CPM values were multiplied by 100 to avoid underflow when estimating model parameters; for brevity, we refer to these as expression values throughout the text.

### 5.4 Targeted subset of genes

Our aim was to focus on eQTL mapping and genomic predictions of expression for a subset of genes of interest; we particularly sought to identify genes that were strong candidates for being regulated by genetic or epigenetic variants. In an earlier study (Crespo-Piazuelo et al. 2023), an eGWAS identified polymorphisms significantly associated (Bonferonni corrected p-values < 0.05) with gene expression that were subsequently grouped into expression quantitative trait locus (eQTL) regions. *Cis*-eQTL regions were defined as those found within a ±1Mb window around their respective gene, and the remaining eQTL regions were denoted as *trans*-eQTLs. Based on these analyses, a set of 7 expressed genes were identified for which top polymorphisms in their respective *cis*-eQTL regions were shared across multiple tissues (duodenum, liver, muscle): *CELF2*, *DET1*, *HUS1*, *NUDT22*, *R3HCC1*, *SLA-7*, and *SUPT3H*. We further identified 3 additional genes that were identified as having *cis*-signals in the eGWAS (Crespo-Piazuelo et al. 2023) and have been previously reported to be methylated: *IGF2*, *PRKAG1*, and *LEPR*, the latter of which was expressed only in the liver. This set of 10 genes was thus used in our subsequent analyses; a description and summary of the previous eGWAS results (Crespo-Piazuelo et al. 2023) for these genes can be found in Supplementary Table 1.

Heritability of tissue-specific expression was estimated by fitting a Bayesian reproducing kernel Hilbert spaces (RKHS) regression (Pérez and de los Campos 2014) using an eigenvalue decomposition of the genomic relatedness matrix based on the centered and scaled genetic variants on the respective chromosome for each target gene (2000 iterations burn-in, followed by 12000 iterations), and calculating the ratio of total additive genetic to total variance. Within-breed learning quality was assessed using the Bayesian RKHS model by calculating the Spearman correlation between estimated and true gene expression values on the learning data for each breed.

### 5.5 Tissue-specific epigenetic annotations

Post-mortem samples were harvested from pig embryos at early organogenesis (*n*=4 pools at 30 days post-fertilisation; dpf) and late organogenesis (*n*=4 individuals; 70 dpf) as well as in newborn LW piglets (*n*=4 individuals; NB). Note that these samples were independent from those used for the whole genome sequencing and transcriptomics assays described above. We focused on epigenetic annotations generated within each of the aforementioned developmental stages in liver and muscle for two different functional genomic assays; these data are indexed and freely available on the GENE-SWitCH project page of the FAANG Data portal (https://data.faang.org/projects/GENE-SWitCH).

First, we considered methylation profiling by whole genome bisulfite sequencing (WGBS) assays for *n*=1 pool or individual at each developmental stage; note that the remaining *n*=3 pools or individuals were profiled by Reduced Representation Bisulfite Sequencing (RRBS) and were not considered here. Genomic regions were categorized with a hidden Markov model (HMM) as being unmethylated (UMR) or lowly methylated (LMR) in a given tissue (Burger et al. 2013; Stadler et al. 2011). Briefly, using the recommended settings for the MethylSeekR package, thresholds for segmentation were as follows: (1) coverage of > 10 reads per site, (2) a *q*-value < 0.05 for regions, and (3) average DNA methylation < 50% for methylome segmentation (de Vos 2023).

Second, chromatin accessibility profiling by assay for transposase-accessible chromatin with high-throughput sequencing (ATAC-seq) was used to identify open chromatin regions (OCR) in *n*=4 animals (2 males and 2 females). ATAC-seq reads were processed with the nf-core ATAC-seq pipeline (https://nf-co.re/atacseq) version 1.2.1. Briefly, reads were mapped to the genome with BWA and open chromatin regions (*i.e.*, broad peaks) were obtained from the mapped reads using Macs2. Peaks from all samples were then merged into 336,746 consensus peaks representing potential regulatory regions, and several quality controls were provided in a multiQC report.

### 5.6 Tissue-agnostic predicted variant effect annotations

The Variant Effect Predictor (VEP) tool (Ensembl release 106, *Sscrofa11.1* reference genome) was used to identify the predicted potential impact of variants (McLaren et al. 2016). Predicted effect classes were collapsed into two broad tissue-agnostic annotations as proposed in (MacLeod et al. 2016): (1) non-synonymous coding (NSC) variants, corresponding to {coding_sequence_variant, frameshift_variant, inframe_deletion, inframe_insertion, initiator_codon_variant, missense_variant, splice_acceptor_variant, splice_donor_variant, splice_region_variant, stop_gained, stop_lost, stop_retained_variant} and (2) potential regulatory (REG) variants, corresponding to {3_prime_UTR_variant, 5_prime_UTR_variant, downstream_gene_variant, mature_miRNA_variant, non_coding_transcript_variant, non_coding_exon_variant, upstream_gene_variant}.

### 5.7 Global set of annotations

Matching tissue-specific epigenetic annotations (OCR, LMR, UMR) at each of the 3 developmental stages (30 dpf, 70 dpf, newborn) for liver and muscle, respectively, were concatenated with tissue-agnostic predicted variant effect categories (REG, NSC). The corresponding genomic intervals were overlapped with SNP positions in the WGS data using the GenomicRanges Bioconductor package (Lawrence et al. 2013). Any remaining unannotated variants were assigned to a final “other” category. The final global set of annotations thus corresponded to a binary matrix with a total of 12 annotation categories in each tissue (10 tissue-specific, 2 tissue-agnostic), with 1’s indicating variants assigned to a particular annotation and 0’s otherwise. To explore similarities among the global set of annotation categories, we performed a hierarchical clustering of annotation categories using complete linkage and the Jaccard distance.

### 5.8 eQTL mapping and prediction with BayesRCπ

A general linear model for genomic prediction can be defined as

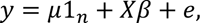

where here *y* is the vector of sex-corrected log_2_ CPM values for a given gene of interest, μ the intercept, *X* the centered and scaled marker matrix, β the vector of SNP effects, *e* the vector of residuals, and *e* follows a centered multivariate normal distribution with variance covariance matrix 𝐼_n_𝜎^2^.

In this work we used chromosome-specific genetic variants to simultaneously perform eQTL mapping and prediction based on each of the three pig breeds considered. In particular, we focus on comparisons between a model based on genomic variants alone with a model additionally including biologically relevant categorizations of variants obtained from complex and overlapping functional annotation maps; to this end, we respectively make use of BayesR (Erbe et al. 2012) and BayesRCπ (Mollandin et al. 2022). Both define prior mixture distributions for β to model variants with null to large effects; BayesRCπ further introduces a mixture of mixtures prior to disambiguate variants with multiple annotations, as is the case in the functional annotation categorizations used here. Given the high resolution of the genomic sequencing data used here, prior mixture distributions for genetic effect sizes were defined for five components, corresponding to 0, 0.001%, 0.01%, 0.1%, and 1% of the total additive genetic variance for both models; otherwise, default parameters were used. Models were fit for each combination of gene (10 target genes), tissue (muscle, liver), and learning breed (DU, LD, LW). Parameters were estimated using a Gibbs sampler for BayesR and BayesRCπ (50,000 iterations, discarding the first 20,000 as burn-in and using a thinning rate of 10).

eQTL mapping was performed using estimates of posterior variances from each model. As genotypes are centered and scaled, for both models the posterior variance of SNP *i* is estimated as 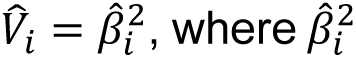 corresponds to the posterior mean of 𝛽_*i*_^2^ across iterations after burn-in. To account for differences in linkage disequilibrium structures across the three breeds, in addition to SNP-specific posterior variances, we also considered window-based posterior variances, where values in non-overlapping windows of 1 Kb were summed rather than considering estimates for individual SNPs; note that one limitation of such an approach is that the number of SNPs summed may not be comparable across windows.

Prediction accuracy across breeds was assessed using the Spearman correlation between true and estimated gene expression values in each of the two breeds not used for learning.

### 5.9 Gene set enrichment analysis

The biological relevance of annotation categories as well as estimated posterior variances from BayesR and BayesRCπ were assessed based on their co-localization with known QTLs in pigs for a variety of pig traits. We made use of the PigQTLdb database (Release 45; *SusScrofa11.1*) to obtain curated cross-experiment results of genotype-phenotype association studies for pig traits (Hu et al. 2021). Only QTLs corresponding to SNPs were considered for the analysis. Gene set enrichment analyses (GSEA) of individual PigQTLdb traits among per-SNP estimated posterior variances were performed using the fgsea Bioconductor package v1.24.0 (Korotkevich et al. 2016). Enrichment tests with an adjusted P-value < 0.05 were considered to be significant. Enrichment analysis results were performed for both individual traits as well as for a set of 5 broad trait categories, the latter of which correspond to the highest level of the PigQTLdb trait hierarchy (Meat and Carcass Traits, Health Traits, Exterior Traits, Production Traits and Reproduction Traits).

### 5.10 Statistical analyses

Principal components analysis (PCA) was performed on genomic data from autosomal chromosomes (chr1 to chr18) using PLINK v1.09 (Chang et al. 2015) and the –pca flag, with per-breed MAF filtering as described above as well as LD-based variant pruning using the -- indep-pairwise flag with a window size of 50 Kb, a step size of 10, and a *r*^2^ threshold of 0.10. PCA was similarly performed on transcriptomic data for each tissue based on uncorrected but scaled log_2_ CPM values using the mixOmics package v 6.28.0 (Rohart et al. 2017). Fisher’s exact test was used to test for the overrepresentation of PigQTLdb QTLs among each annotation category. Unless otherwise noted, statistical analyses were performed using R v4.4.0.

## Acknowledgments

Computing cluster resources were provided by the Centre de Traitement de l’Information Génétique (CTIG) of the INRAE Animal Genetics department. We are grateful to Juan Pablo Sánchez for helpful discussions about this work. The authors thank the selection companies Axiom and Nucléus of Alliance R&D and the INRAE UE3P France Génétique Porc facility (https://doi.org/10.15454/1.5573932732039927E12) for producing the LW pigs, as well as Selección Batallé S.A. for producing the DU pigs and Hendrix Genetics for producing the LD pigs. The authors report there are no competing interests to declare.

## CRediT authorship contribution statement

**Fanny Mollandin**: Formal analysis, Investigation, Methodology, Software, Visualization, Writing – original draft, Writing – review and editing. **Hervé Acloque**: Funding acquisition, Resources, Writing – review and editing. **Maria Ballester**: Investigation, Writing – review and editing. **Marco Bink**: Supervision, Writing – review and editing. **Mario Calus**: Supervision, Writing – review and editing. **Daniel Crespo-Piazuelo**: Data curation, Formal analysis, Investigation, Writing – review and editing. **Pascal Croiseau**: Methodology, Supervision, Writing – review and editing. **Sarah Djebali**: Formal analysis, Writing – review and editing. **Sylvain Foissac**: Data curation, Formal analysis, Writing – review and editing. **Hélène Gilbert**: Supervision, Writing – review and editing. **Elisabetta Giuffra**: Funding acquisition, Writing – review and editing. **Cervin Guyomar**: Formal analysis, Writing – review and editing **Ole Madsen**: Supervision, Writing – review and editing. **Marie-José Mercat**: Writing – review and editing. **Bruno da Costa Perez**: Supervision, Writing – review and editing. **Jani de Vos**: Data curation, Formal analysis, Writing – review and editing. **Andrea Rau**: Conceptualization, Formal analysis, Investigation, Methodology, Supervision, Visualization, Writing – original draft, Writing – review and editing.

## Data availability

Raw sequence data supporting the findings of this study are available at the FAANG data portal (https://data.faang.org), with the following BioProject accession codes: WGS data (<u>PRJEB58030</u>), RNA-seq data (<u>PRJEB58031</u>), WGBS data (<u>PRJEB42772</u>) and ATAC-seq data (<u>PRJEB44468</u>). The BayesR and BayesRCπ models are available as part of the open-source BayesRCO package, available at https://github.com/FAANG/BayesRCO. Source code to reproduce the analyses and manuscript figures and full results from the GSEA of PigQTLdb traits among BayesR and BayesRCO eQTLs are available at https://github.com/andreamrau/2025_BayesRCO_eQTL, with associated raw data available at https://doi.org/10.5281/zenodo.14973198.

## Declaration of generative AI and AI-assisted technologies in the writing process

During the preparation of this work the corresponding author used ChatGPT in order to write the abstract of this manuscript. After using this tool, the authors reviewed and edited the content as needed and take full responsibility for the content of the publication.

