## Supplementary Tables and Figures for "Guiding eQTL mapping and genomic prediction of gene expression in three pig breeds with tissue-specific epigenetic annotations from early development"

#### Outline

Supplementary Tables ........................................ 1
Supplementary Figures ........................................ 6

#### Supplementary Tables

**Supplementary Table 1**: Description of genes used in the study, including Ensembl ID, gene name, chromosome and position, and number of *cis*- and *trans*-eQTLs in liver and muscle identified from (Crespo-Piazuelo et al. 2023). *Cis*-eQTLs were defined as variants located within ±1 Mb of the gene. *Trans*-eQTLs were defined as the sum of variants located more than ±1 Mb on the same chromosome as the gene and those located on other chromosomes. As *LEPR* has liver-specfic expression, no values are included for it in muscle.

| Ensembl ID | Gene | Chr | Pos | *Cis* (liver) | *Trans* (liver) | *Cis* (muscle) | *Trans* (muscle) |
|---|---|---|---|---|---|---|---|
| ENSSSCG00000005103 | *DET1* | 1 | 191,263,447-191,284,968 | 3 | 124+1 | 3 | 177+4 |
| ENSSSCG00000013039 | *NUDT22* | 2 | 7,880,258-7,883,945 | 217 | 0+6 | 295 | 27+3 |
| ENSSSCG00000035293 | *IGF2* | 2 | 1,469,132-1,496,442 | 18 | 0+1 | 354 | 17+79 |
| ENSSSCG00000000185 | *PRKAG1* | 5 | 15,033,053-15,049,660 | 488 | 0+5 | 0 | 2+2 |
| ENSSSCG00000025188 | *LEPR* | 6 | 146,798,979-146,896,108 | 342 | 2+16 | — | — |
| ENSSSCG00000001398 | *SLA-7* | 7 | 23,634,639-23,649,314 | 9 | 53+1 | 13 | 44+1 |
| ENSSSCG00000001709 | *SUPT3H* | 7 | 39,751,927-40,161,114 | 25 | 0+0 | 247 | 9+10 |
| ENSSSCG00000011121 | *CELF2* | 10 | 60,506,482-61,084,861 | 4 | 0+0 | 5 | 12+6 |
| ENSSSCG00000039915 | *R3HCC1* | 14 | 7,405,792-7,434,935 | 1348 | 54+2 | 2947 | 42+7 |
| ENSSSCG00000028523 | *HUS1* | 18 | 48,522,997-48,538,788 | 0 | 0+0 | 7443 | 70+1 |

**Supplementary Table 2.** Number of variants before and after filtering, and number of annotated variants each category (after merging developmental stages) per chromosome.

| Chr | # variants before filtering | # variants after filtering | Annotated variants, liver | | | Annotated variants, muscle | | | VEP |
|---|---|---|---|---|---|---|---|---|---|
| | | | OCR | UMR | LMR | OCR | UMR | LMR | |
| 1 | 1,815,427 | 629,069 | 17019 | 20959 | 34972 | 16421 | 15816 | 21425 | 41536 |
| 2 | 1,264,180 | 535,945 | 16816 | 15225 | 36076 | 17467 | 11841 | 18921 | 49002 |
| 5 | 905,184 | 305,552 | 10745 | 10482 | 17460 | 9823 | 6829 | 11782 | 27919 |
| 6 | 1,466,687 | 514,990 | 20344 | 16057 | 22847 | 18682 | 11216 | 14877 | 40812 |
| 7 | 1,075,681 | 426,627 | 16671 | 13448 | 22214 | 13474 | 9533 | 13707 | 45388 |
| 10 | 903,373 | 341,017 | 10536 | 10922 | 25887 | 11859 | 7867 | 14561 | 27365 |
| 14 | 1,247,911 | 480,183 | 15987 | 11855 | 24404 | 15022 | 8526 | 14945 | 28072 |
| 18 | 542,356 | 177,240 | 5075 | 5447 | 7956 | 4059 | 3735 | 5111 | 12239 |
| Total | 9,220,799 | 3,410,623 | 113,193 | 104,395 | 191,816 | 106,807 | 75,363 | 115,329 | 272,333 |

**Supplementary Table 3.** Significant (adjusted P-value < 5%) GSEA enrichments for reported PigQTLdb QTL trait categories based on estimated posterior variances according to BayesRCπ with annotations. ES: enrichment score; NES: normalized enrichment score.

| Broad QTL category | QTL category | Adj p-value | Log2 error | ES | NES | Size | Gene | Tissue | Breed |
|---|---|---|---|---|---|---|---|---|---|
| Health | LDL cholesterol | 0.012 | 0.498 | 0.907 | 1.226 | 45 | IGF2 | liver | LD |
| Meat and carcass | Oleic acid content | 0.016 | 0.432 | 0.835 | 1.069 | 326 | R3HCC1 | muscle | LD |
| Meat and carcass | Palmitic acid content | 0.016 | 0.432 | 0.836 | 1.071 | 325 | R3HCC1 | muscle | LD |
| Meat and carcass | Palmitoleic acid content | 0.016 | 0.455 | 0.835 | 1.069 | 355 | R3HCC1 | muscle | LD |
| Health | White blood cell number | 0.016 | 0.477 | 0.901 | 1.157 | 71 | R3HCC1 | muscle | LD |
| Meat and carcass | Monounsaturated fatty acid content | 0.020 | 0.432 | 0.834 | 1.069 | 327 | R3HCC1 | muscle | LD |
| Meat and carcass | Saturated fatty acid content | 0.021 | 0.407 | 0.830 | 1.063 | 371 | R3HCC1 | muscle | LD |
| Meat and carcass | Backfat at tenth rib | 0.021 | 0.477 | 0.941 | 1.198 | 29 | LEPR | liver | LW |
| Meat and carcass | Stearic acid content | 0.023 | 0.407 | 0.829 | 1.061 | 377 | R3HCC1 | muscle | LD |
| Meat and carcass | Subcutaneous fat thickness | 0.031 | 0.432 | 0.861 | 1.129 | 75 | PRKAG1 | muscle | LD |
| Meat and carcass | Monounsaturated fatty acid content | 0.034 | 0.407 | 0.797 | 1.087 | 327 | R3HCC1 | liver | LD |
| Meat and carcass | Oleic acid content | 0.034 | 0.432 | 0.797 | 1.087 | 326 | R3HCC1 | liver | LD |
| Meat and carcass | Palmitic acid content | 0.034 | 0.455 | 0.803 | 1.094 | 325 | R3HCC1 | liver | LD |
| Meat and carcass | Palmitoleic acid content | 0.034 | 0.407 | 0.796 | 1.086 | 355 | R3HCC1 | liver | LD |
| Health | Red blood cell count | 0.045 | 0.381 | 0.935 | 1.245 | 19 | R3HCC1 | muscle | LD |
| Meat and carcass | Shear force | 0.045 | 0.455 | 0.897 | 1.214 | 35 | NUDT22 | liver | LD |
| Meat and carcass | Lean meat percentage | 0.046 | 0.432 | 0.923 | 1.252 | 24 | NUDT22 | liver | LD |
| Meat and carcass | Saturated fatty acid content | 0.049 | 0.381 | 0.791 | 1.078 | 371 | R3HCC1 | liver | LD |
| Meat and carcass | Stearic acid content | 0.049 | 0.381 | 0.791 | 1.078 | 377 | R3HCC1 | liver | LD |

**Supplementary Table 4.** Significant (adjusted P-value < 5%) GSEA enrichments for reported PigQTLdb QTL trait categories based on estimated posterior variances according to BayesR (genomic data alone). ES: enrichment score; NES: normalized enrichment score.

| Broad QTL category | QTL category | Adj p-value | Log2 error | ES | NES | Size | Gene | Tissue | Breed |
|---|---|---|---|---|---|---|---|---|---|
| Health | Mean corpuscular hemoglobin concentration | 0.000 | 0.627 | 0.942 | 2.180 | 10 | DET1 | liver | LW |
| Reproduction | Teat number | 0.006 | 0.477 | 0.792 | 1.863 | 32 | PRKAG1 | liver | LW |
| Meat and carcass | Obesity index | 0.010 | 0.498 | 0.817 | 2.092 | 12 | SUPT3H | liver | LW |
| Meat and carcass | Oleic acid content | 0.014 | 0.477 | 0.906 | 2.062 | 10 | NUDT22 | liver | LW |
| Exterior | Cannon bone circumference | 0.030 | 0.432 | 0.759 | 1.427 | 42 | SUPT3H | muscle | DU |
| Production | Femur length | 0.030 | 0.432 | 0.854 | 1.576 | 21 | SUPT3H | muscle | DU |
| Production | Humerus length | 0.030 | 0.455 | 0.846 | 1.566 | 24 | SUPT3H | muscle | DU |
| Production | Ulna length | 0.030 | 0.432 | 0.856 | 1.576 | 20 | SUPT3H | muscle | DU |
| Meat and carcass | Linoleic acid content | 0.034 | 0.432 | 0.858 | 2.070 | 14 | NUDT22 | liver | LW |
| Meat and carcass | Monounsaturated fatty acid to polyunsaturated fatty acid ratio | 0.041 | 0.407 | 0.858 | 2.090 | 15 | NUDT22 | liver | LW |
| Meat and carcass | Polyunsaturated fatty acid content | 0.041 | 0.407 | 0.855 | 2.084 | 15 | NUDT22 | liver | LW |
| Production | Body length | 0.041 | 0.407 | 0.806 | 1.493 | 22 | SUPT3H | muscle | DU |
| Production | Tibia length | 0.041 | 0.407 | 0.844 | 1.565 | 25 | SUPT3H | muscle | DU |
| Meat and carcass | Monounsaturated fatty acid content | 0.043 | 0.407 | 0.439 | 1.437 | 327 | R3HCC1 | liver | DU |
| Meat and carcass | Oleic acid content | 0.043 | 0.407 | 0.441 | 1.442 | 326 | R3HCC1 | liver | DU |
| Meat and carcass | Palmitic acid content | 0.043 | 0.407 | 0.442 | 1.447 | 325 | R3HCC1 | liver | DU |
| Meat and carcass | Palmitoleic acid content | 0.043 | 0.381 | 0.435 | 1.426 | 355 | R3HCC1 | liver | DU |
| Meat and carcass | Saturated fatty acid content | 0.043 | 0.407 | 0.436 | 1.427 | 371 | R3HCC1 | liver | DU |
| Meat and carcass | Stearic acid content | 0.043 | 0.381 | 0.428 | 1.400 | 377 | R3HCC1 | liver | DU |
| Health | Mean corpuscular volume | 0.047 | 0.455 | 0.653 | 1.888 | 112 | SLA-7 | muscle | LD |

**Supplementary Table 5**: Per-breed Pearson correlations of standardized expression values with the first five principal components (PC) based on the top 80% of variants according to estimated BayesRCπ posterior variances (Supplementary Figure 8). Cases considered include one of weak prediction portability(*SUPT3H* expression in muscle, with learning on DU) and strong prediction portability (*HUS1* expression in muscle, with learning on DU).

| PC | SUPT3H muscle (DU learning) | | | HUS1 muscle (DU learning) | | |
|---|---|---|---|---|---|---|
| | DU | LD | LW | DU | LD | LW |
| 1 | -0.215 | -0.038 | 0.165 | -0.015 | -0.024 | 0.078 |
| 2 | -0.076 | 0.244 | 0.156 | 0.240 | -0.136 | 0.050 |
| 3 | -0.244 | -0.139 | 0.097 | 0.148 | 0.048 | 0.050 |
| 4 | -0.274 | -0.098 | -0.153 | -0.171 | 0.044 | 0.003 |
| 5 | 0.169 | 0.051 | -0.082 | 0.166 | 0.172 | -0.130 |

### Supplementary Figures

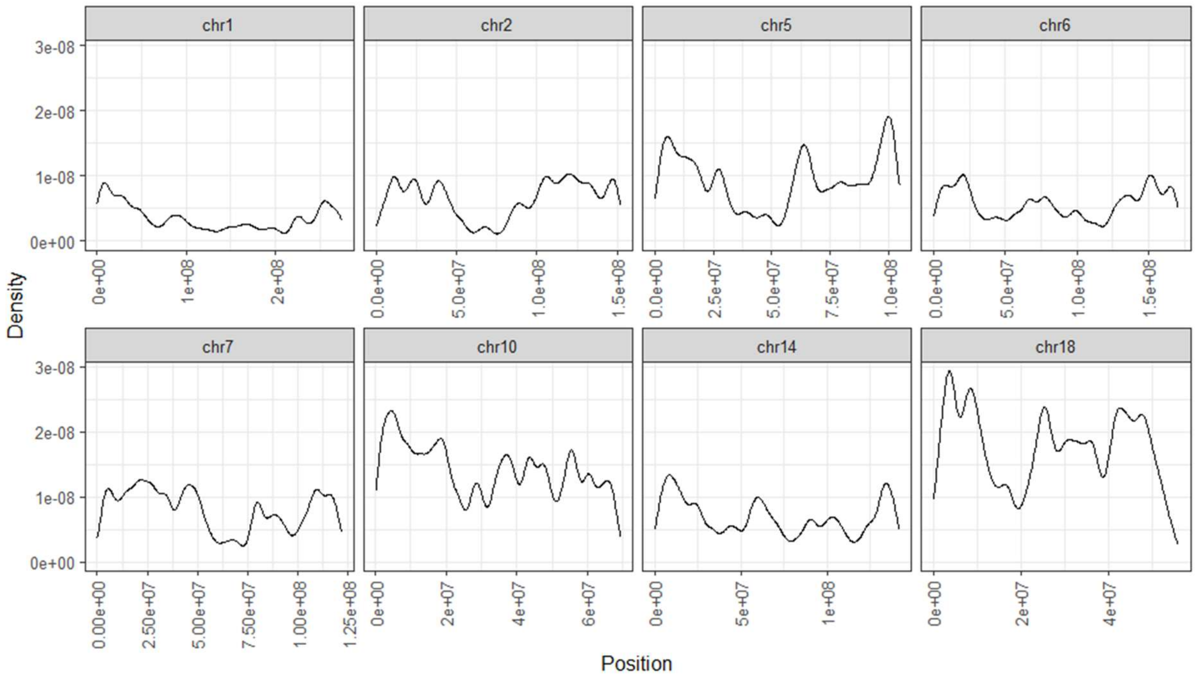


**Supplementary Figure 1.** Density plots with respect to the position of filtered variants for each chromosome.

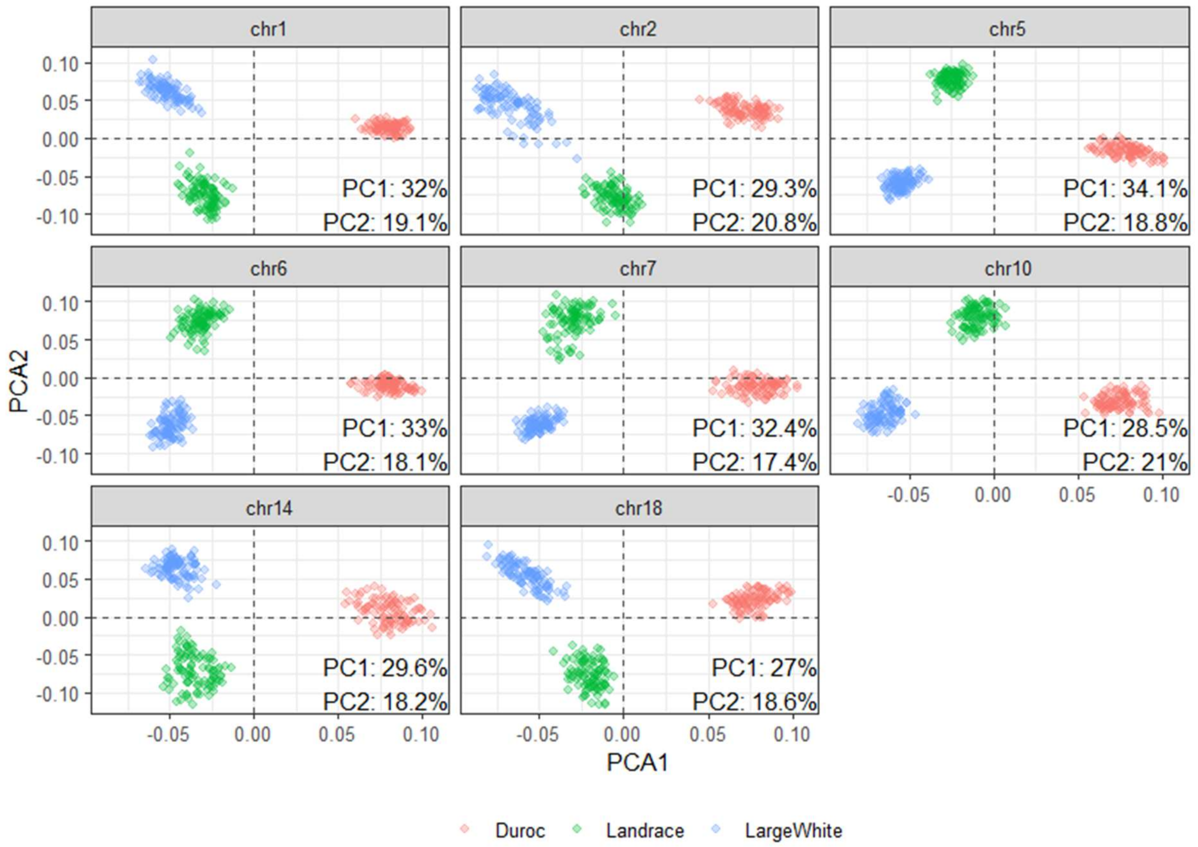


**Supplementary Figure 2.** Plots of first two principal components (PC) from chromosome-specific genomic PCA, with breeds indicated by color (red, Duroc; green, Landrace; blue Large White). Percentage variance explained for each of the first two PCs is indicated in text for each chromosome.

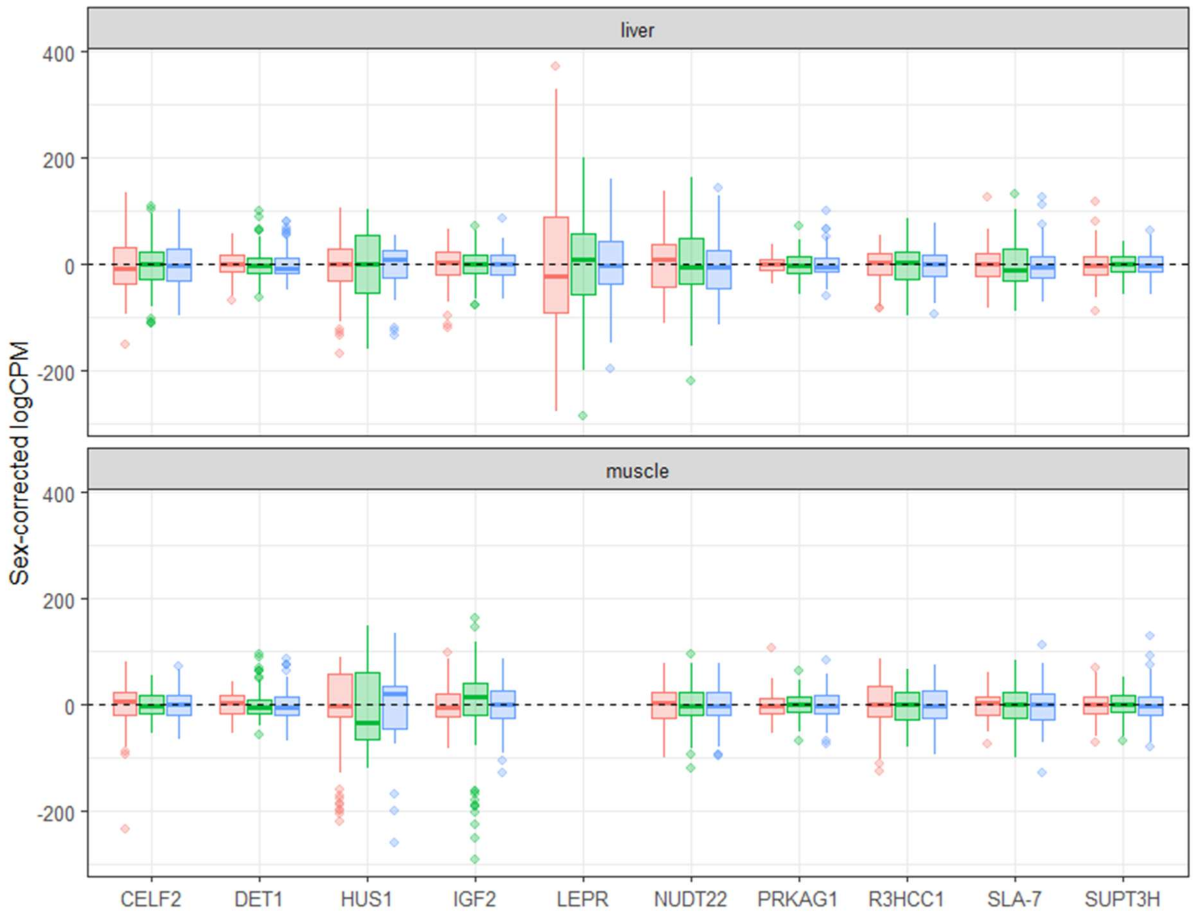


**Supplementary Figure 3.** Distribution of sex-corrected log-CPM values in liver (top) and muscle (bottom) for target gene list, colored by breed (red, Duroc; green, Landrace; blue Large White).

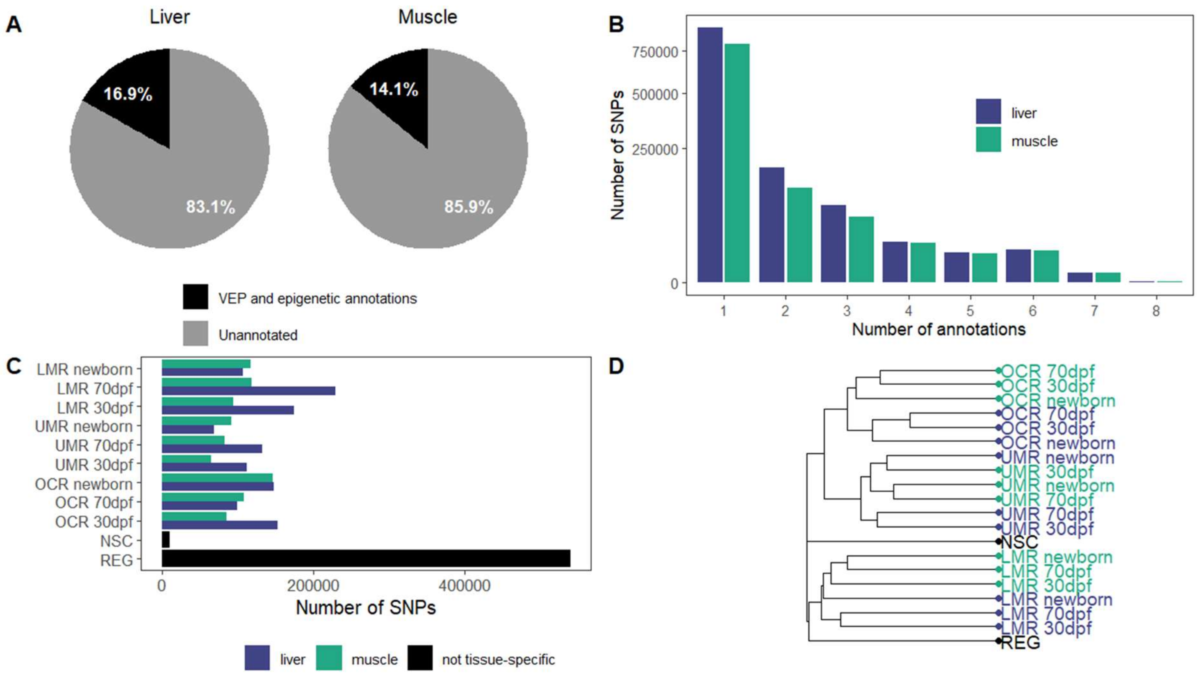


**Supplementary Figure 4.** (A) Percentage of SNPs annotated according to tissue-agnostic variant effect predictions (VEP) or tissue-specific epigenetic annotations in liver (left) and muscle (right). (B) Number of SNPs annotated with one or more annotation categories in liver (blue) and muscle (green). (C) Number of annotated SNPs for each category. (D) Dendrogram representing a hierarchical clustering of annotation categories based on the Jaccard distance and complete linkage. Liver- and muscle-specific epigenetic annotations are represented in blue and green, respectively, and VEP annotations in black. OCR: open chromatin regions; UMR: unmethylated regions: LMR: lowly methylated regions; REG: predicted regulatory variant; NSC: predicted nonsynonymous coding variants; dpf: days post fertilization.

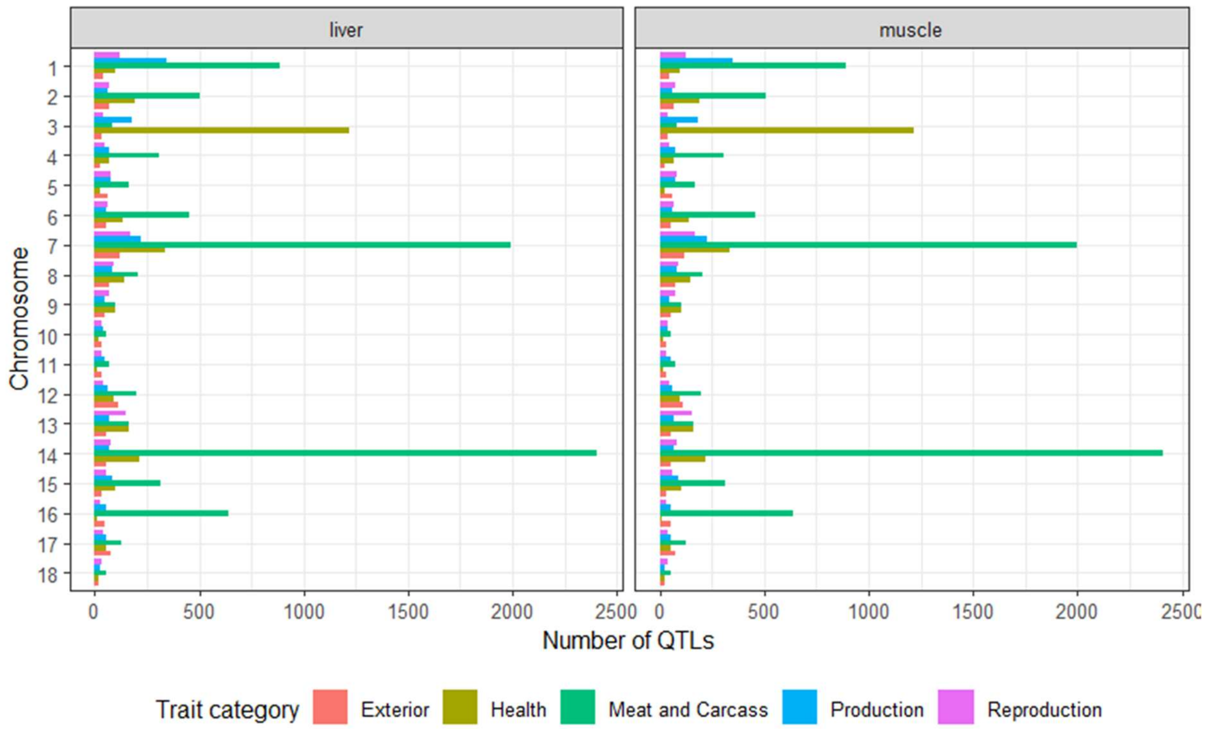


**Supplementary Figure 5.** Frequency of QTLs for each chromosome in liver (left) and muscle (right) corresponding to broad trait categories from PigQTLdb.

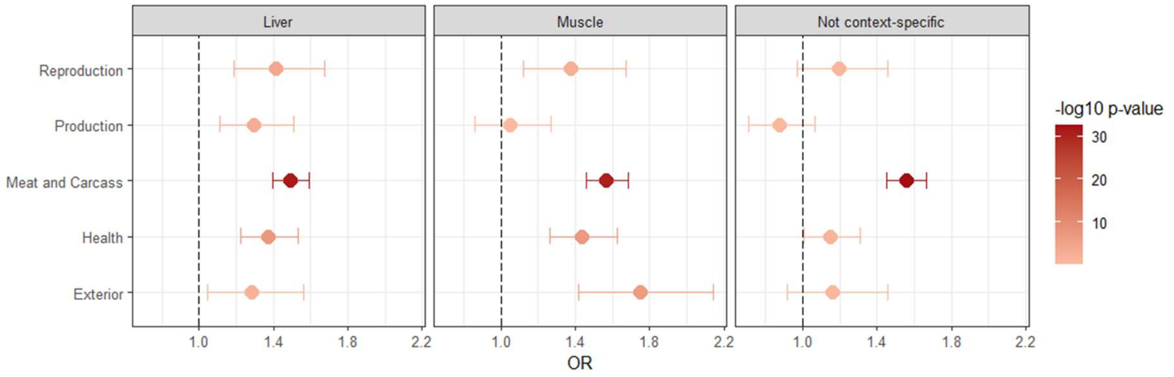


**Supplementary Figure 6.** Odds Ratio (OR) and corresponding -$\log_{10}$ P-values from a Fisher's exact test of the enrichment of liver-specific epigenetic marks (left; UMR, LMR and OCR from 30 dpf, 70 dpf and newborn stages), muscle-specific epigenetic marks (middle; UMR, LMR and OCR from 30 dpf, 70 dpf and newborn stages), and non tissue-specific variant effect predictions (right) for each broad trait category from PigQTLdb. The dotted reference line corresponds to an OR of 1 (i.e., no enrichment).

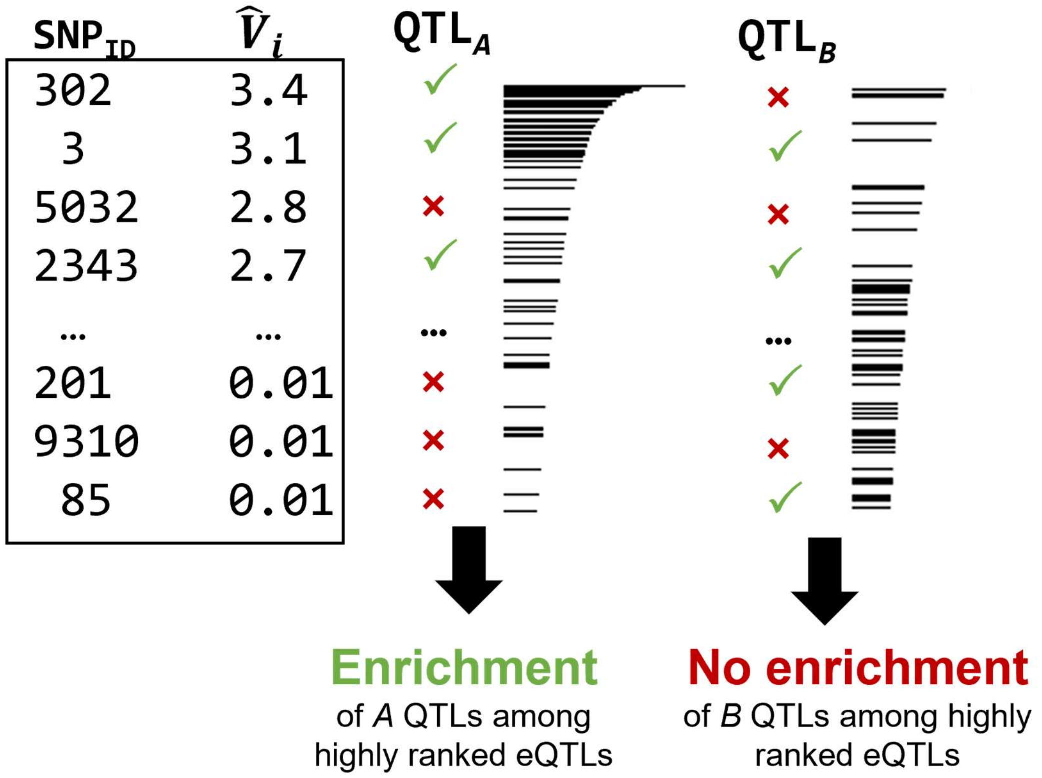


**Supplementary Figure 7**: Schematic illustration of the enrichment analyses of performance QTLs among highly ranked eQTLs based on BayesRCπ posterior variances.

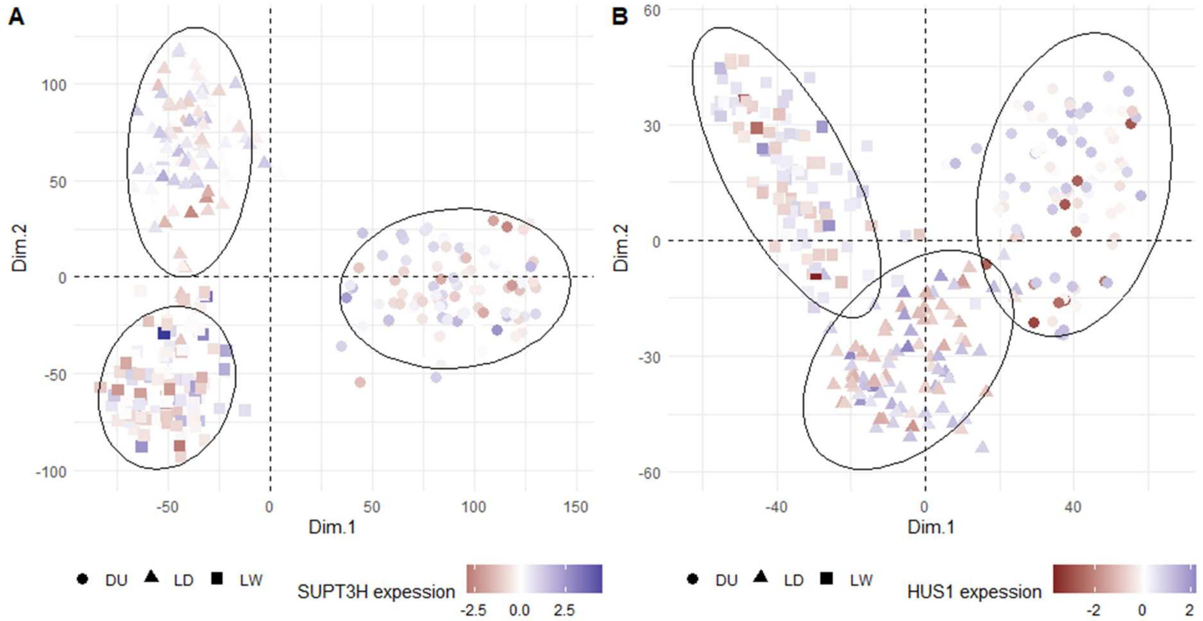


**Supplementary Figure 8**. Plots of first two principal components (PC) from the top 80% of variants according to estimated BayesRCπ posterior variances. Breeds are indicated with shapes (DU: circles, LD: triangles, LW: squares) and corresponding ellipses, and points are colored according to standardized gene expression. (A) A case of weak prediction portability: *SUPT3H* expression in muscle, with learning on DU. (B) A case of strong prediction portability: *HUS1* expression in muscle, with learning on DU.

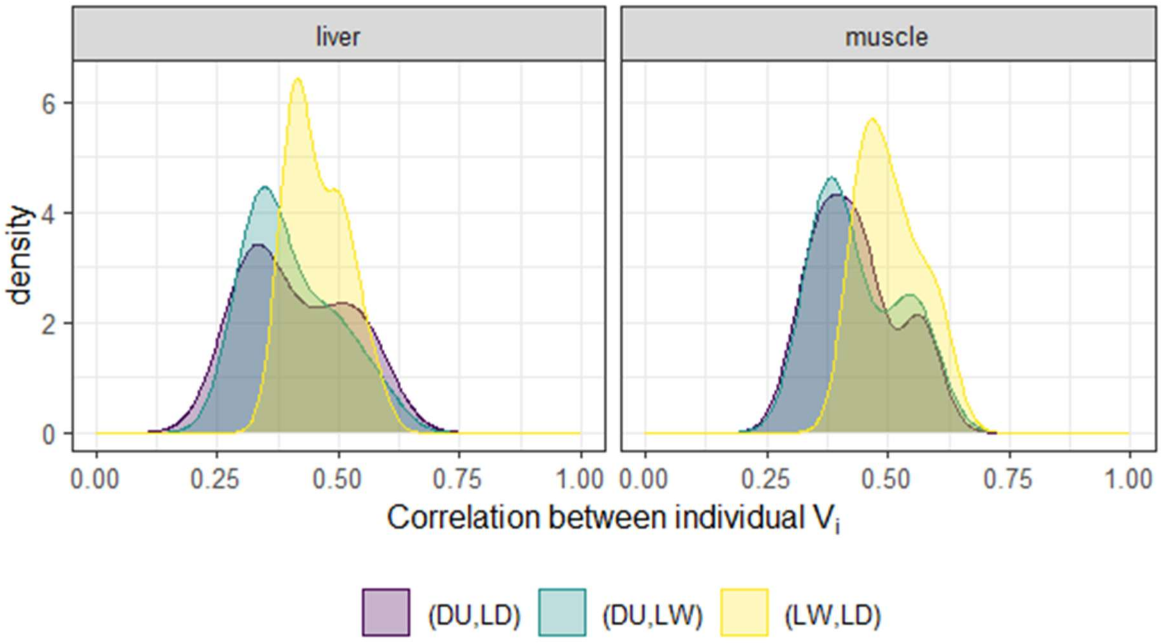


**Supplementary Figure 9**. Distribution of the pairwise correlations of per-variant estimated posterior variances between pairs of breeds in liver (left) and muscle (muscle) across genes. Each pairwise combination of breeds is represented by a distinct color.

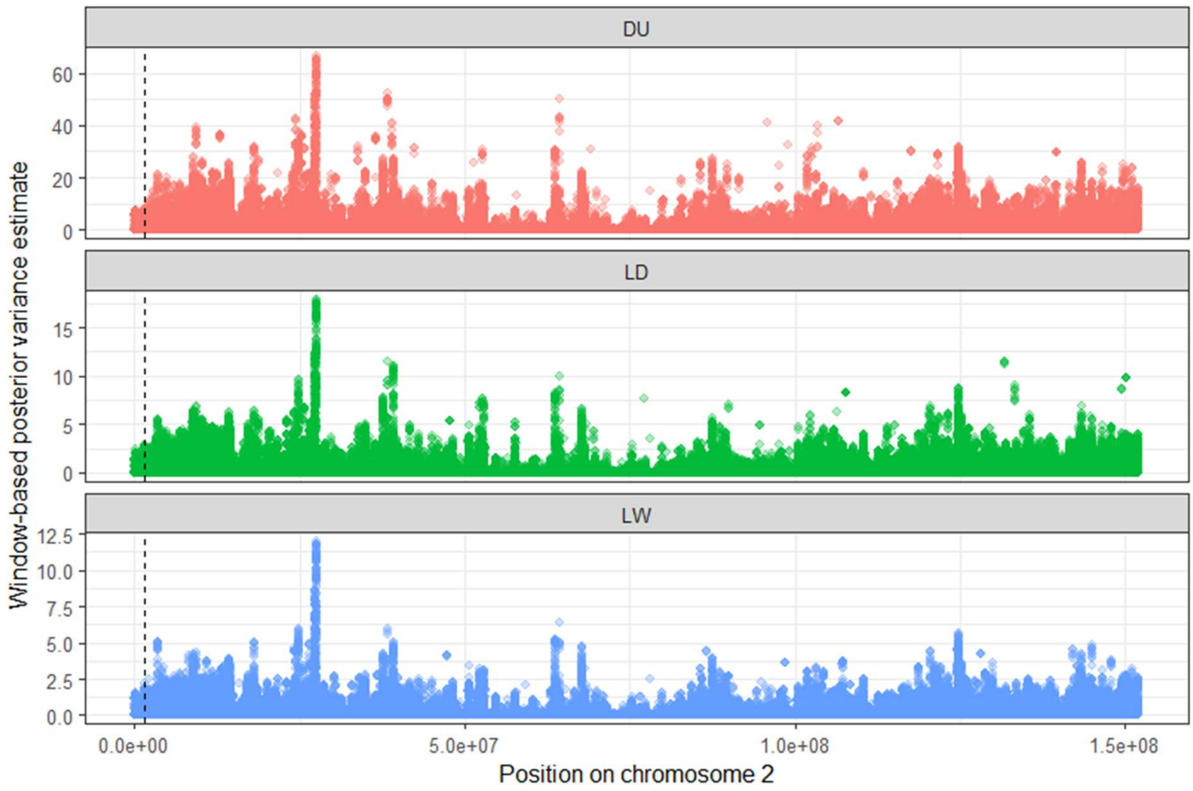


**Supplementary Figure 10**: Manhattan plot of summed window-based posterior variances for *IGF2* (chromosome 2) for each of the 3 learning breeds in liver. The dotted vertical line indicates the position of *IGF2*.

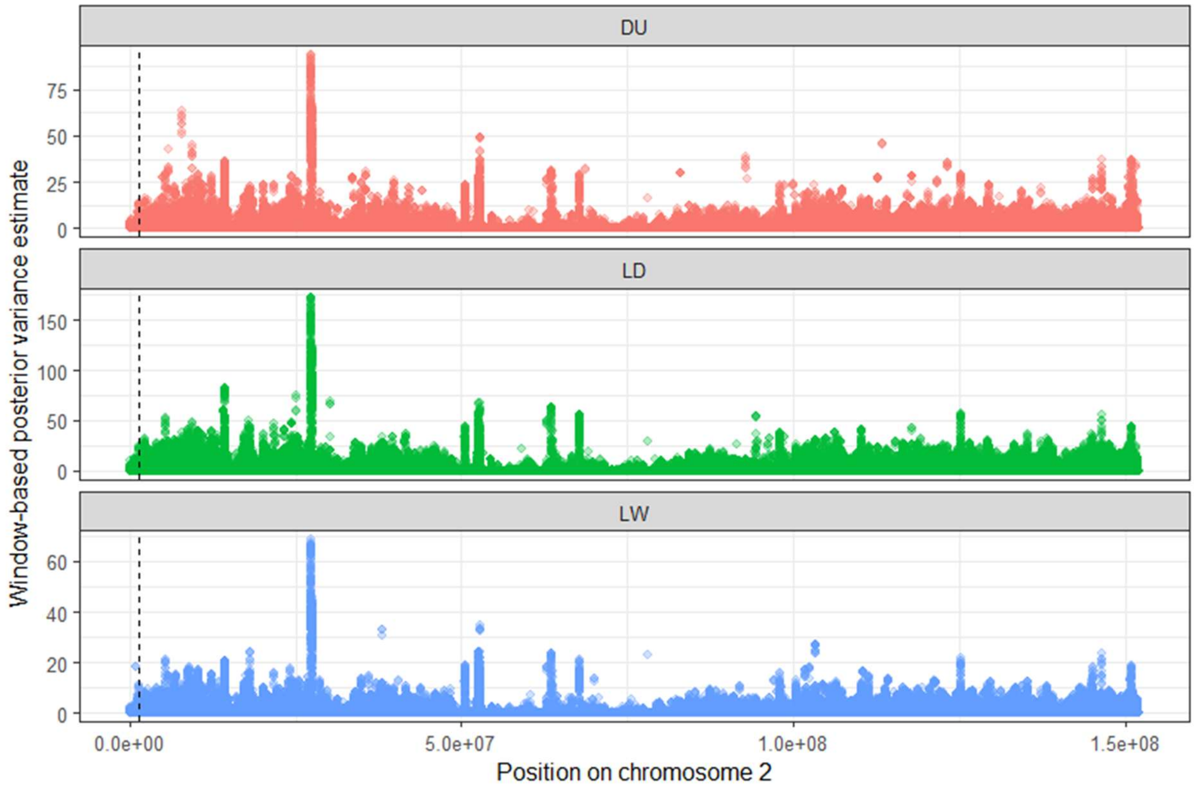


**Supplementary Figure 11**: Manhattan plot of summed window-based posterior variances for *IGF2* (chromosome 2) for each of the 3 learning breeds in muscle. The dotted vertical line indicates the position of *IGF2*.

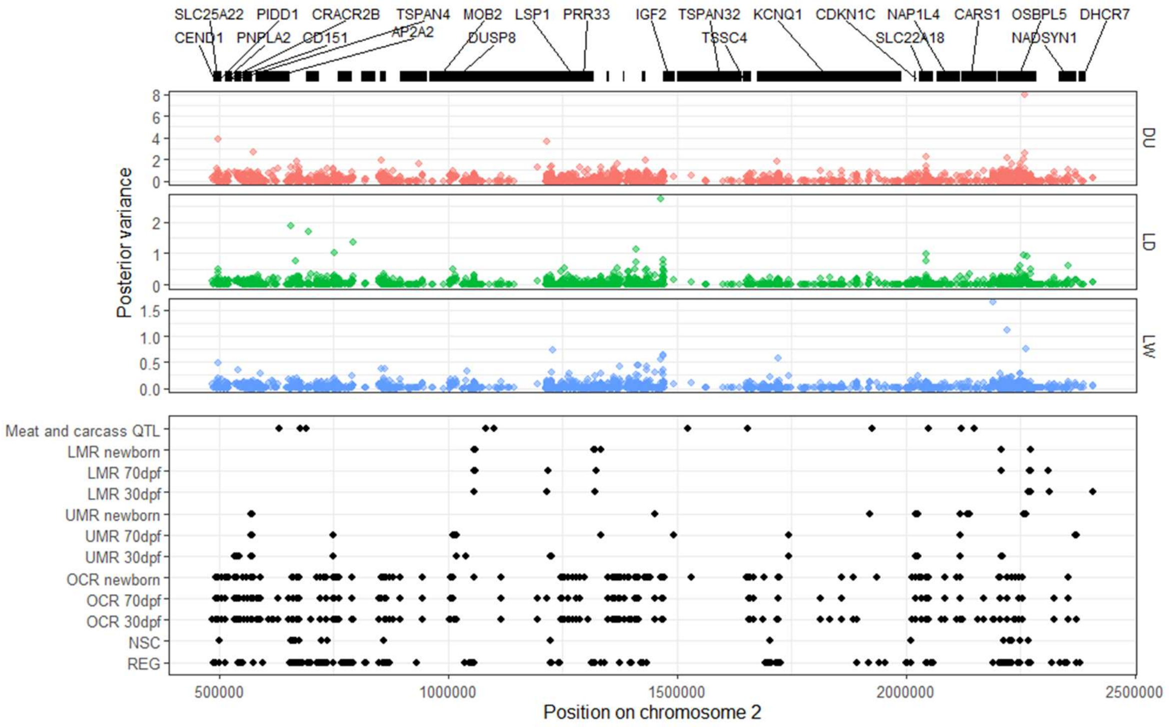


**Supplementary Figure 12**: (A) Neighborhood +/- 1 Mb around *IGF2*. Gene positions are highlighted in the top panel with black bars. Estimated posterior variances of effects for each genetic variant in the window are shown in the middle panel for each learning breed (DU=Duroc: LD = Landrace; LW = Large White). The bottom panel indicates the position of predicted variant effect annotations, liver-specific epigenetic annotations, and QTL categories from PigQTLdb.

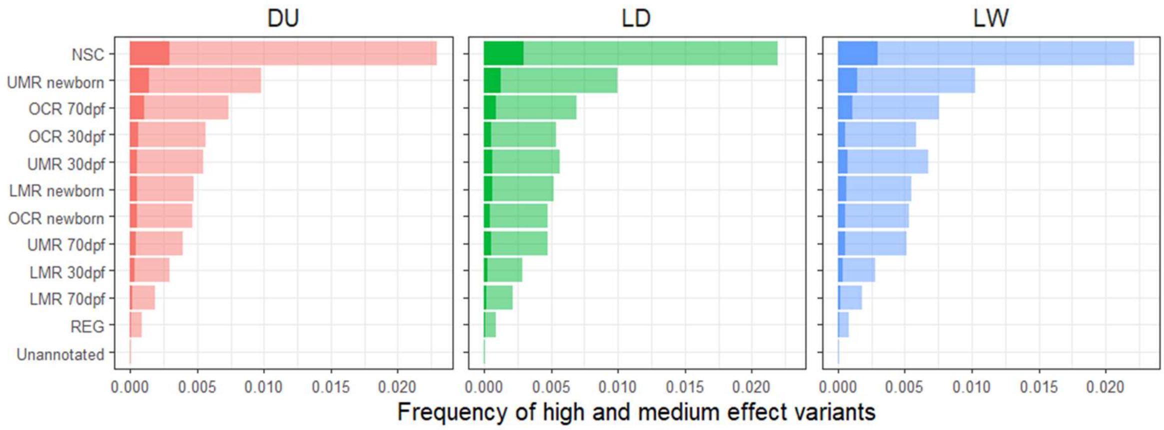


**Supplementary Figure 13.** BayesRCπ frequency of VEP and liver-specific epigenetic annotations among medium (lighter shading) or high (darker shading) effect SNPs for each of the three learning breeds.
